# A Risk-reward Examination of Sample Multiplexing Reagents for Single Cell RNA-Seq

**DOI:** 10.1101/2023.06.20.544880

**Authors:** Daniel V. Brown, Casey J.A. Anttila, Ling Ling, Patrick Grave, Tracey M. Baldwin, Ryan Munnings, Anthony J. Farchione, Vanessa L. Bryant, Amelia Dunstone, Christine Biben, Samir Taoudi, Tom S. Weber, Shalin H. Naik, Anthony Hadla, Holly E. Barker, Cassandra J. Vandenberg, Genevieve Dall, Clare L. Scott, Zachery Moore, James R. Whittle, Saskia Freytag, Sarah A. Best, Anthony T. Papenfuss, Sam W.Z. Olechnowicz, Sarah E. MacRaild, Stephen Wilcox, Peter F. Hickey, Daniela Amann-Zalcenstein, Rory Bowden

## Abstract

Single-cell RNA sequencing (scRNA-Seq) has emerged as a powerful tool for understanding cellular heterogeneity and function. However the choice of sample multiplexing reagents can impact data quality and experimental outcomes. In this study, we compared various multiplexing reagents, including MULTI-Seq, Hashtag antibody, and CellPlex, across diverse sample types such as human peripheral blood mononuclear cells (PBMCs), mouse embryonic brain and patient-derived xenografts (PDXs). We found that all multiplexing reagents worked well in cell types robust to *ex vivo* manipulation but suffered from signal-to-noise issues in more delicate sample types. We compared multiple demultiplexing algorithms which differed in performance depending on data quality. We find that minor improvements to laboratory workflows such as titration and rapid processing are critical to optimal performance. We also compared the performance of fixed scRNA-Seq kits and highlight the advantages of the Parse Biosciences kit for fragile samples. Highly multiplexed scRNA-Seq experiments require more sequencing resources, therefore we evaluated CRISPR-based destruction of non-informative genes to enhance sequencing value. Our comprehensive analysis provides insights into the selection of appropriate sample multiplexing reagents and protocols for scRNASeq experiments, facilitating more accurate and cost-effective studies.

## Introduction

Single-cell RNA sequencing (scRNA-Seq) has been powered by advancements in molecular biology, microfluidics, and high-throughput sequencing (1). Applications of scRNA-Seq span cell atlases, pooled screens and clinical studies (2). Single-cell approaches have expanded to include additional modalities, such as surface protein measurement, open chromatin analysis, and CRISPR perturbation (3). As scRNA-Seq becomes more accessible, increased sample sizes and biological replicates enhance scientific rigor but necessitate larger, more complex experiments. Although the cost per cell is decreasing, overall experimental costs remain high.

Batch effects, which encompass technical variation introduced during sample preparation, library preparation, and sequencing, contribute significantly to the complexity of singlecell data analysis (4). These effects demand increased analyst time and can potentially attenuate biological signals.

Sample multiplexing has emerged as an elegant solution to these challenges. The concept was first demonstrated by mixing genetically distinct samples and subsequently deconvoluting them using genotypes called in the sequencing data (5). However, in many studies the absence of natural genetic variation renders this approach unfeasible. As an alternative, various methods have been developed to deliver exogenous sample-identifying DNA barcodes. The first implementation utilized oligo-tagged antibodies targeting ubiquitous cell surface proteins (6). Subsequent technologies have delivered DNA barcodes via lipids, concanavalin A, click chemistry, transfection, or transduction (7–11).

Regardless of the delivery mechanism, sample multiplexing necessitates additional upfront handling of individual samples, with the potential to perturb cell states (12) or reduce viability. Antibody- and lipid-based barcodes have become the most popular systems, for their broad applicability and ease of use. Recently, Mylka et al., directly compared these multiplexing methods, recommending different solutions for different sample types (13).

The advent of commercial fixed scRNA-Seq kits, such as the Parse Biosciences Evercode kits and 10x Genomics Flex, has effectively decoupled sample collection from processing. Both kits incorporate sample multiplexing into their molecular biology workflows, eliminating the need for specific labeling steps. The Parse Biosciences Evercode kits utilize multiwell plates, enabling sample multiplexing by dispensing each sample into a separate well. In contrast, the 10x Genomics Flex kit employs a ligation probe-based assay, where sample barcodes are embedded in the probe sequences.

We have conducted an extensive comparison of antibody, lipid-based, and fixed sample multiplexing reagents across diverse and broadly representative cell types, including human peripheral blood mononuclear cells (PBMCs), mouse embryonic brain, and ovarian carcinosarcoma patient-derived xenografts (PDX). We evaluate CRISPR-based destruction of non-informative genes, an important potential adjunct in controlling the cost of larger single-cell experiments. Through upfront optimization and downstream comparative analyses, we propose guidelines for experimental design and the utilization of different protocols in various contexts.

## Results

### Comparison of multiplexing reagents in human PBMCs

To evaluate the performance of sample multiplexing reagents (Table S1) in a system where we could obtain ground truth from SNP genotypes, we analyzed PBMCs from four human donors (Figure 1A).

**Fig. 1.**
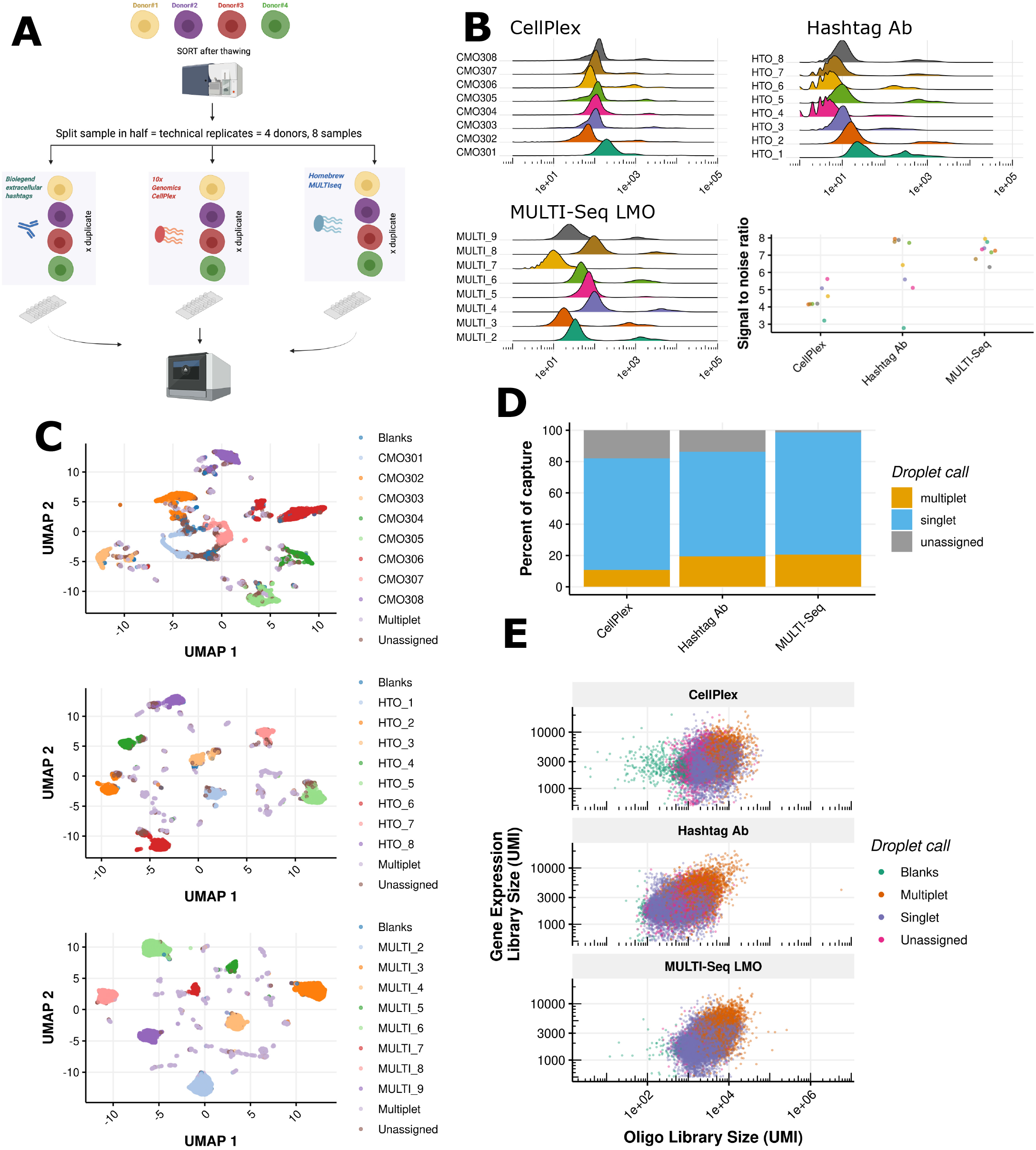
Benchmarking sample multiplexing reagents in human PBMCs. (A) Experimental design. PBMCs isolated from four unrelated healthy donors were divided into technical duplicate prior to labeling. Each protocol was captured in a separate 10x Genomics v3.1 reaction. (B) Log10 transformed oligo tag counts for each multiplexing protocol with a summary of signal-to-noise. Signal-to-noise is defined as the difference between the mean background (left) and foreground (right) oligo tag counts on a log scale divided by variance. (C) UMAP dimension reduction visualisation of multiplexing oligo tag counts for each protocol. Cells are coloured by Cell Ranger multi call. (D) Summary of multiplexing tag calls per protocol as reported by Cell Ranger multi. Blanks and unassigned are both reported as unassigned. (E) Relationship between oligo tag and gene expression library size for each protocol tested.

We first undertook a round of optimization by flow cytometry, titrating the Total-Seq hashtag antibody to a concentration of ten-fold less than the manufacturer’s recommendations (0.1 *μ*g per reaction) (Figure S1A). We next substituted the polyA capture sequence of the described MULTI-Seq oligo (7) with the 10x Genomics feature barcode 2 sequence (Figure S1B). Titrating the MULTI-Seq lipid modified oligos (LMO) resulted in a rapid loss of signal, therefore we used a concentration of 200 nM as reported in the original study (Figure S1C).

Each PBMC donor sample was divided into technical duplicates for the sample multiplexing labeling reaction (Figure 1A) and captured with 10x Genomics v3.1 chemistry at a cell input of 35,000 cells, for a theoretical output of 20,000 cell-containing droplets at a 16.11% doublet rate (Satija lab calculator).

After library preparation and sequencing, we examined the count distributions for each tag and protocol (Figure 1B). CellPlex had the lowest signal-to-noise and highest proportion of unassigned cells (Figure 1C and D). Of note the doublet rate is higher for hashtag antibody and MULTI-Seq than the theoretical 16.11% expected from loading each 10x Genomics capture with 35,000 cells. We later titrated the CellPlex reagent ten-fold below the manufacturer’s recommendations without a loss in signal (Figure S3).

The signal-to-noise was most consistent for MULTI-Seq with hashtag antibody also performing well aside from a single labeling reaction, HTO_1, which had a high background and lower signal indicating an issue with the antibody reagent rather than an error in sample handling. The HTO_1 sample’s poor signal to noise negatively impacted other tags. Dimension reduction of the hashtag antibody capture revealed satellite clusters with a subset of cells having a correct dominant tag but contaminated by HTO_1 (Figure S2B).

Examination of the relationship between oligo tag library and gene expression library size revealed multiplets had a higher library size than singlets (Figure 1E). In contrast, unassigned cells had a similar library size to singlets, suggestive of a failure in oligo tag labeling rather than unassigned cells being enriched from empty droplets or damaged cells.

### Accuracy of sample multiplexing oligos compared to SNP genotypes

We next compared the accuracy of the sample multiplexing tag assignments to ground truth SNP assignments (Figure 2A). Given our experimental design with four unrelated donors in duplicate, the doublet rate was lower from SNP calls (12.08%) than from multiplexing tags (16.56%). The identifiable doublet rate excludes homotypic doublets.

**Fig. 2.**
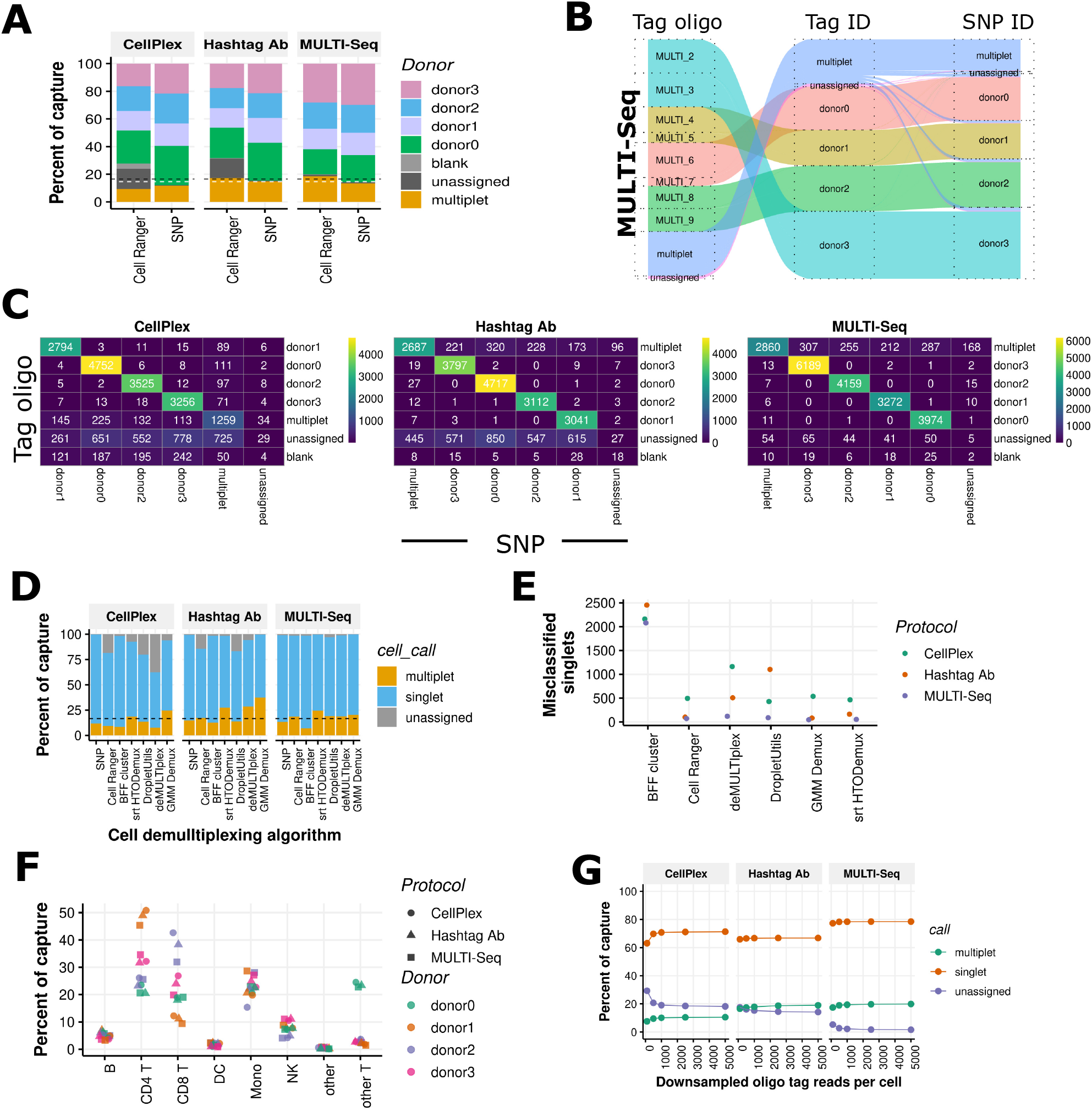
Comparison of cell demultiplexing software. (A) Comparison of oligo tag calls with Cell Ranger multi versus SNP calls. The white horizontal dashed line represents the theoretical doublet rate based on 4 SNP donors.The black line represents doublet rate based on eight multiplexed samples. (B) Alluvial plot for MULTI-Seq LMO comparing demultiplexing on a tag basis (left), donor basis (middle) and corresponding SNP calls (right). (C) Heatmap of the sample multiplexing oligo-snp assignment contingency table. (D) Comparison of droplet calls from multiple demultiplexing algorithms implemented in cellhashR. (E) Donor misclassification rate by algorithm, based on SNP ground truth. Unassigned droplets and multiplets were removed. (F) Immune subset composition by donor and protocol. Automated annotation is shown with Level 1 granularity. (G) Downsampling analysis of oligo tag libraries. Oligo tag libraries were downsampled to a fixed number of reads per cell and each dataset was demultiplexed with Cell Ranger multi.

The proportions of individual donors recovered when comparing SNPs to sample multiplexing oligos was similar aside from the unassigned category (Figure 2A). This was confirmed upon closer inspection of the multiplexing tag calls to SNP calls in an alluvial plot and heatmap (Figure 2B and C). The major discordant droplets were non-identifiable doublets when calling multiplets based on 4 SNPs versus 8 multiplexing oligos. Having used the Cell Ranger multi algorithm for sample demultiplexing, we next compared other algorithms available in the cellhashR package (14). The three sample multiplexing datasets generated had different characteristics with CellPlex having lower signal to noise, hashtag antibody having one poorly performing tag and MULTI-Seq being a high quality dataset (Figure 2D).

Cell Ranger multi performed as well as other algorithms in hashtag antibody and MULTI-Seq datasets, despite a warning during runtime that these multiplexing tag oligos are not supported. In the CellPlex dataset, the Bimodal Flexible Fitting (BFF cluster) algorithm performed best. In contrast, in the high-quality MULTI-Seq dataset, BFF Cluster called four-fold more false positives, assigning more multiplets to singlets (Figure 2D). We computed the overall classification accuracy, (OCA) the same metric used in Mylka et al., (13), (Table 1). The OCA is defined as the sum of matching assignments between demultiplexing tags and SNP assignments divided by the count of all cell-containing droplets. Consistent with the signal-to-noise metrics, MULTI-Seq performed best, followed by hashtag antibody with CellPlex the poorest performing protocol.

**Table 1.** Overall classification accuracy. OCA is the number of matching assignments between sample multiplexing tags and SNP assignments divided by all cellcontaining droplets.

| Protocol | OCA | Unassigned |
| --- | --- | --- |
| MULTI-Seq | 0.926 | 0.0153 |
| Hashtag Ab | 0.804 | 0.145 |
| CellPlex | 0.761 | 0.185 |

To evaluate the impact of sequencing depth on demultiplexing accuracy, we downsampled the multiplexing tag library sequencing data and reprocessed the data using Cell Ranger Multi. The recommended number of reads per cell by 10x Genomics is 5,000 for hashtag antibodies and CellPlex. Beyond 1,000 reads per cell, an increase in the number of reads per cell exerted a negligible effect on demultiplexing performance (Figure 2G). An alternate guideline provided on the 10x Genomics website indicates 1,000 usable oligo tag reads per cell, which is consistent with our findings. Particularly for high-quality datasets, a total of 5,000 reads per cell for sample multiplexing oligos appears excessive.

### Comparison of sample multiplexing reagents in mouse embryonic brain

PBMCs are a robust sample type that do not require dissociation and can be maintained as a single-cell suspension for a prolonged period on ice with only minor effects on viability or phenotype (15). We next aimed to benchmark sample multiplexing reagents in mouse embryonic brain E18.5, a more challenging tissue (Figure 3A).

**Fig. 3.**
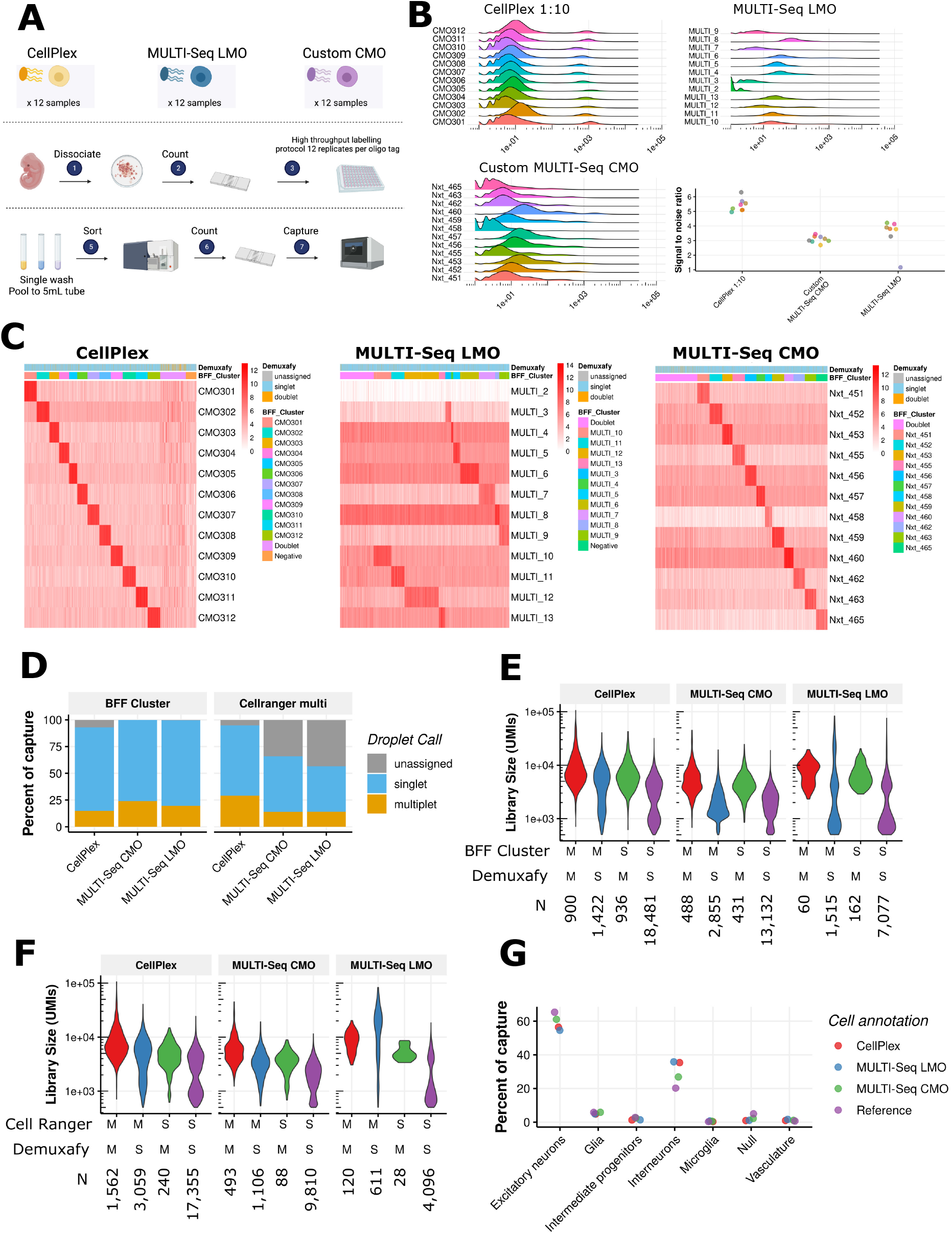
Benchmarking sample multiplexing reagents in mouse embryonic brain. (A) Experimental design. Embryonic day 18.5 mouse brain from a single animal was used. The single cell suspension was split into 12 partitions for each reagent tested. A high throughput labeling protocol in 96 well plates was used. Cells were pooled prior to FACS sorting. (B) Log10 transformed oligo tag counts for each multiplexing protocol with a summary of signal-to-noise. (C) Heatmaps of oligo tag counts annotated by BFF Cluster call and Demuxafy consensus call. (D) Identity of cell containing droplets from BFF Cluster and Cell Ranger multi algorithms. (E) Library size of gene expression library based on Cell Ranger demultiplexing calls. (F) Library size of gene expression library based on BFF Cluster demultiplexing calls. (G) Cell annotations recovered from each protocol. Seurat map query level 1 granularity is shown.

In contrast to PBMCs, which were processed using a lowthroughput labeling protocol in 1.5mL tubes, we utilized a high-throughput labeling protocol in a 96 well plate. We also adjusted the FACS sort step as per 10x Genomics guidelines to conduct the sort after labeling with multiplexing oligos, rather than sorting after thawing and before labeling as in the PBMC experiment.

However, in a pilot experiment, we were unable to detect any signal using mouse hashtag antibodies with this sample type (Figure S5A). Thus, as a substitute for the hashtag antibody we evaluated a cholesterol modified custom MULTISeq oligo, composed of the CellPlex oligo sequence grafted onto the MULTI-Seq lipid (Figure S5B).

Evaluation of the sequencing counts of the multiplexing tag libraries revealed a good separation of signal and background for diluted CellPlex (Figure 3B). In contrast the two MULTISeq designs had poorer signal to background. During laboratory processing there was incomplete removal of supernatants due to concern over loss of cell pellets. With a cell input of 100,000 per well, pellets were invisible. The remaining dis-sociation media may have inhibited MULTI-Seq LMO, as it is quenched by proteins (7). In line with this observation, MULTI-Seq CMO which is not quenched by culture media performed comparatively better (Figure 3C).

A further logistical issue related to shifting the sorting step until after multiplexing oligo labeling to conform to 10x Genomics supported protocols. This change necessitated a sequential sort of sample pools in the order MULTI-Seq LMO, MULTI-Seq CMO and CellPlex. Viability measurements dropped from 93% to 76% live cells between MULTI-Seq LMO and CellPlex for this reason (Figure S5H). The number of cell-containing droplets retrieved was consistent with a drop in viability over time (Table S2).

Since CellPlex performed the best in this experiment and had the shortest time between cell sorting and single-cell capture, we later assessed if prolonged storage on ice had any effect on signal-to-noise metrics (Figure S6). Indeed 30 minutes storage on ice increased background and reduced signal for the lipid based CellPlex reagent.

In the mouse embryo experiment, the MULTI-Seq reagents yielded low signal-to-noise ratios and a high proportion of unassigned droplets (Figure 3D). As BFF cluster performed well in the low quality CellPlex PBMC dataset, we also assessed its performance in the E18.5 mouse embryo cells. Indeed, the proportion of droplets assigned to multiplets and singlet samples also increased.

As a single sample from an inbred mouse strain was used for this experiment, we were unable to use SNP genotypes as a ground truth. We therefore utilized a doublet detection software package, Demuxafy, based on gene expression as an alternative source of droplet identity information. Demuxafy is a wrapper program around many common algorithms (16). Comparison of droplet calls from demultiplexing algorithms based on multiplexing tags with calls based on gene expression (Figure 3E) showed little difference in gene expression library size between concordant multiplets and singlets. More singlets were recovered from the overlap of Demuxafy and BFF Cluster compared to the number recovered from the overlap of Demuxafy and Cell Ranger Multi.

### Comparison of sample multiplexing reagents in human tumour nuclei

Having evaluated sample multiplexing reagents in intact cells, we next compared the reagents in single-nucleus preparations. Nuclei more faithfully represent cell type composition in tissues that are difficult to dissociate enzymatically than whole-cell preparations (17, 18). Here, we used human ovarian carcinosarcoma patientderived xenograft (PDX) tissue that was snap frozen as tissue pieces according to 10x Genomics best practices. Due to the differing growth rates of the different PDX models it was impractical to process all samples fresh on the same day. We prepared nuclei with a 10x Genomics isolation kit and immediately performed a capture of unlabelled nuclei to identify the effect of prolonged sample handling required by labeling steps on the quality of the transcriptome. We subsequently labelled the remaining nuclei with three different multiplexing reagents (Figure 4A).

**Fig. 4.**
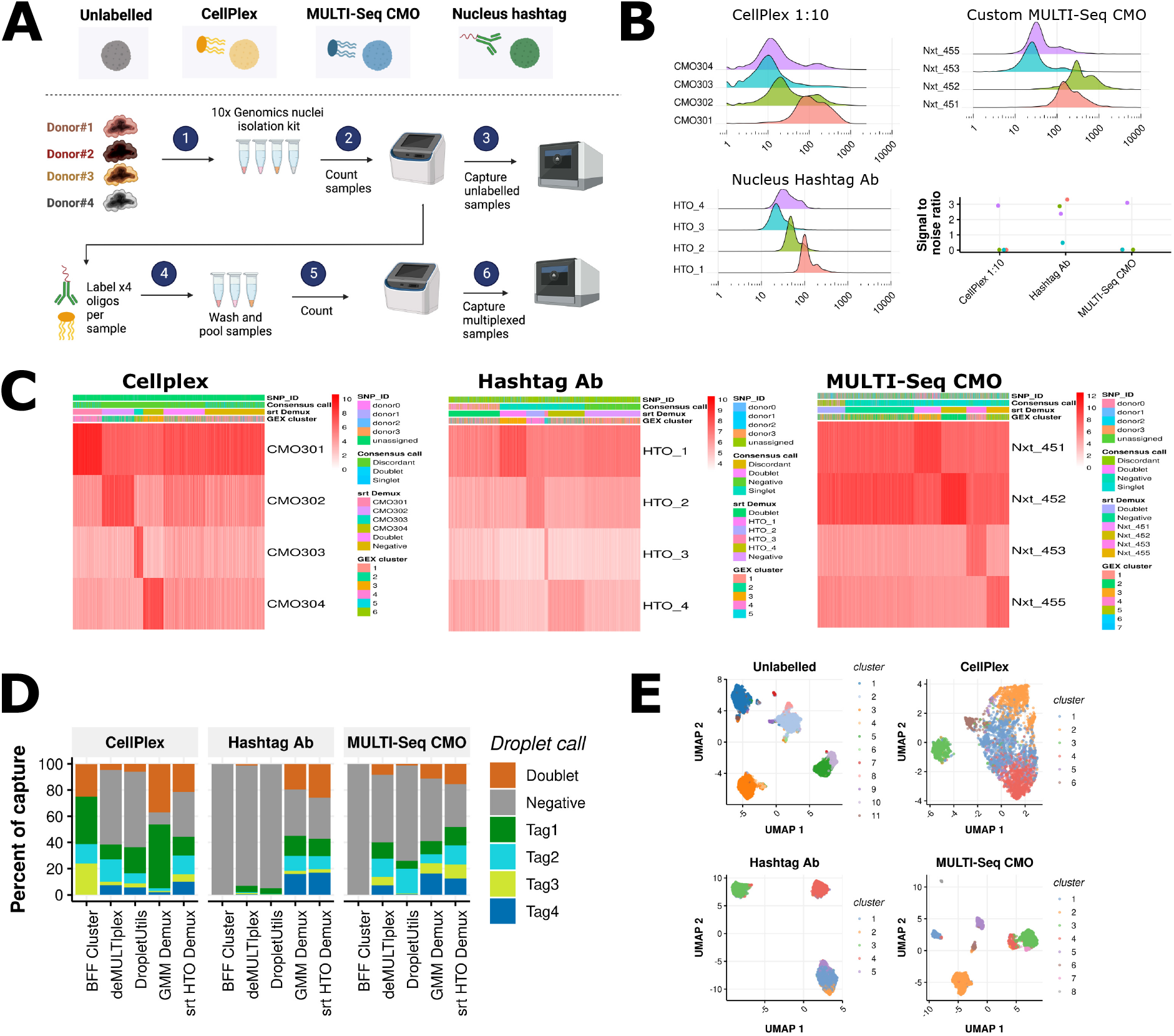
Benchmarking sample multiplexing reagents in ovarian carcinosarcoma xenograft nuclei. (A) Experimental design. Nuclei were isolated from four PDX donor samples, performing an immediate microfluidics capture, followed by labeling the remaining nuclei with multiplexing oligos. (B) Log10-transformed oligo tag counts for each multiplexing protocol with a summary of signal-to-noise ratios. (C) Heatmaps of oligo tag counts annotated by cellhashR consensus call, Seurat HTODemux call, and cluster identity from parallel gene expression data. Cells (columns) are ordered by HTODemux call. (D) Comparison of droplet calls made by demultiplexing algorithms. Calls were generated with cellhashR. (E) UMAP dimension reduction visualization of gene expression data for each protocol. Cells are colored by gene expression cluster identity.

The quality of the nuclei preparation immediately after isolation was good but visually deteriorated after the multiplexing oligo labeling step (Figure S7A). The CellPlex sample experienced a clog resulting in a wetting failure and low recovery volume. While Souporcell estimated an ambient RNA content of 29.97% for droplets from the unlabelled sample, it could not generate an estimate for samples labelled with sample multiplexing oligos. In the labelled samples, the majority of cells could not be assigned to a SNP donor, likely due to a low molecular complexity and fraction of reads in cells (Figure S7B).

The corresponding signal-to-noise ratio of the oligo tag libraries was poor with low separation from background (Figure 4B). Cell ranger multi failed to assign the majority of nuclei to samples (Figure 4C), (Figure S7D). We used the cellhashR package to compare droplet assignment methodologies; Seurat (srt) HTODemux performed the best under the theoretical expectation that samples would be in equal proportions (Figure 4D).

Since each microfluidic capture contained cells from four PDX models, we next checked the gene expression data for separation by sample of origin after dimension reduction (Figure 4E). The unlabelled sample showed four major clusters reflecting the SNP donors with minor satellite clusters reflecting cell doublets. Overlaying these gene expression cluster labels with the oligo tag calls provided confirmation HTODemux calls were largely sample specific (Figure S7E).

### Evaluation of fixed single-nucleus RNA-Seq in human tumour nuclei

In the light of the poor performance of sample multiplexing oligos in nuclei from solid tumour samples, we evaluated fixed snRNA-Seq kits from Parse Biosciences (mini Evercode v2) and 10x Genomics (Flex v1, 4 barcodes) (Figure 5A). Immediately after fixation the single nucleus suspension appeared free of debris and clumps (Figure S8A). Following probe hybridization and sample pooling multiple washes are required for 10x Genomics Flex. With each centrifugation and resuspension step, the sample became increasingly more clumpy (Figure S8B), resulting in a clogged microfluidics chip and failed capture. For the Parse Biosciences experiment, 2 of the 4 samples suffered excessive clumping and were omitted (Figure S8D).

**Fig. 5.**
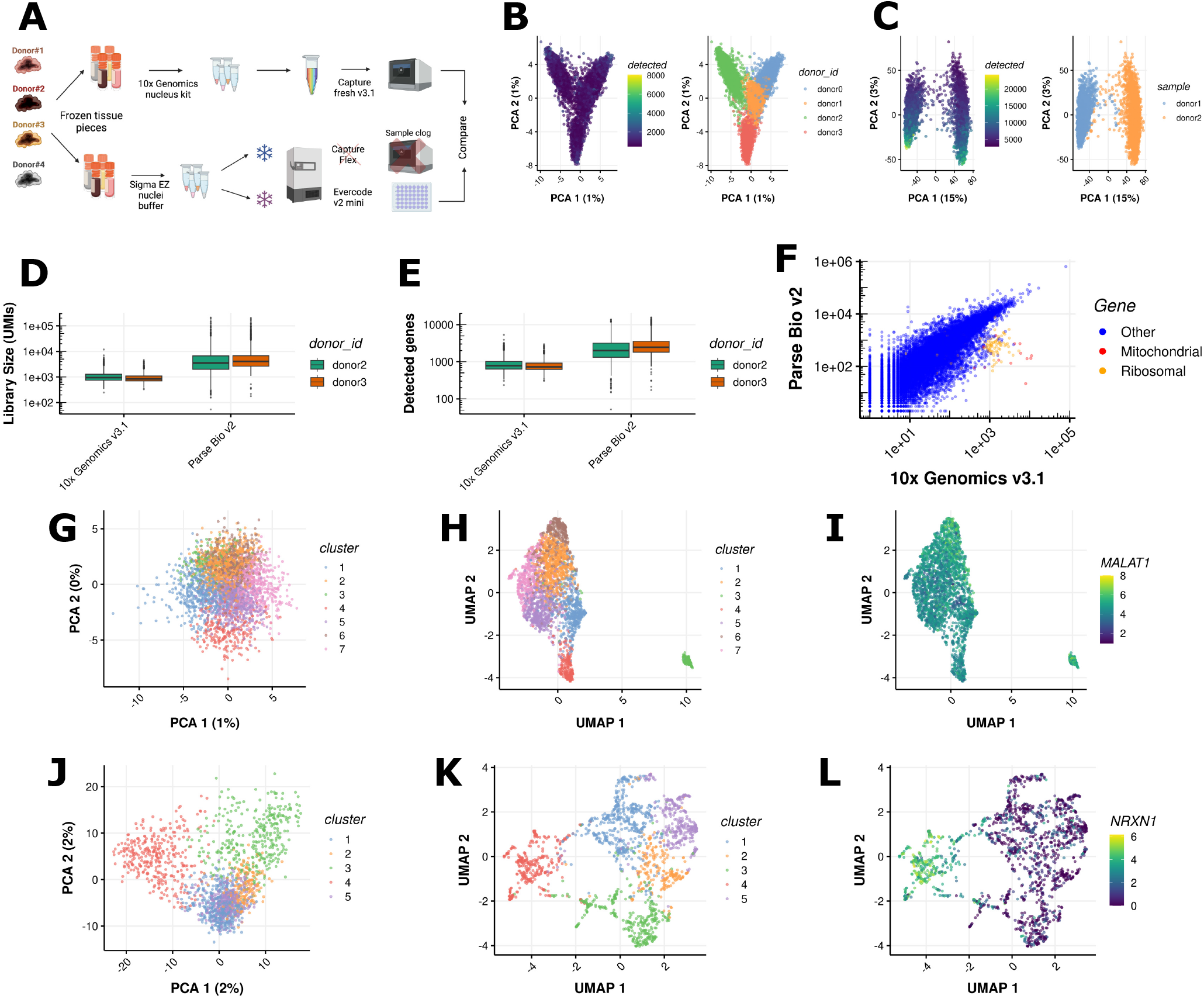
Benchmarking fixed scRNA-Seq in Ovarian carcinosarcoma xenografts. (A) Experimental design. (B) PCA of 10x Genomics v3.1 fresh nuclei prior to sample demultiplexing. Cells coloured by number of detected genes and SNP donor or origin.Percentage of variance explained by the first two principal components is indicated. (C) PCA of Pase Biosciences v2 fixed nuclei prior to sample demultiplexing. Cells coloured by number of detected genes and reverse transcription barcode.. (D) Library size comparison between scRNA-Seq protocols. Sequencing reads were downsampled to an equivalent number per cell. (E) Gene detection comparison between protocols. (F) Gene expression comparison between scRNA-Seq protocols, each dot is the sum of counts across all single cells for a gene. (G) PCA of 10x Genomics v3.1 data for PDX donor 1 coloured by cluster. (H) UMAP of 10x Genomics v3.1 data for PDX donor 1. (I) PCA of 10x Genomics v3.1 data for PDX donor 1 coloured by *MALAT1* expression. (J) PCA of Parse Biosciences v2 data for PDX donor 1 coloured by cluster. (K) UMAP of Parse Biosciences v2 data for PDX donor 1. (L) UMAP of Parse Biosciences v2 data for PDX donor 1 coloured by *NRXN1* expression.

Since the 10x Genomics Flex capture was unsuccessful, we compared Parse Biosciences data to the unlabelled 10x Genomics v3.1 fresh nucleus dataset. Parse Biosciences had more reads in cells (Table S4) reflecting a more efficient use of sequencing resources. Accordingly, the library sizes and number of detected genes were approximately five fold higher in Parse Biosciences when downsampling to an equivalent number of reads per library (Figure 5D and E). Consistent with the manufacturer’s specifications, the cell doublet rate was an order of magnitude lower for Parse Biosciences. There was a high concordance (Pearson correlation 0.918) in gene expression between technologies (Figure 5F). Given that the sample were nuclei, the relatively high mitochondrial transcript content in the 10x Genomics v3.1 data was unexpected.

Importantly, the Parse Biosciences v2 dataset exhibited greater biological variation compared to the 10x Genomics v3.1 dataset. In the latter, an outlier cluster containing a high proportion of the lncRNA *MALAT1* distorted the dimension reduction results (Figure 5H). These cells persisted even after excluding cells with low library sizes and high mitochondrial gene percentages. In contrast, the Parse Biosciences dataset captured biologically meaningful variation, revealing a distinct cluster expressing the *NRXN1* gene (Figure 5I and Figure S8F). Neurexin-1-alpha is a cell adhesion protein and may represent a more epithelial-like subpopulation within the tumor (19, 20).

### Evaluation of CRISPRclean destruction of abundant genes

Multiplexing tags reduce per-cell costs for cell capture and library preparation by increasing the yield of singlecell partitioning. However, superloading does not decrease the cost of sequencing. Instead, it can increase required sequencing volumes due to higher doublet rates and the need to sequence the sample multiplexing tag library. Thus methods that can reduce sequencing requirements are arguably of enhanced value for multiplexed experiments.

We evaluated the Jumpcode Genomics CRISPRclean Single Cell RNA Boost Kit in reducing the amount of uninformative sequence data by using a guide RNA library targeting unaligned reads, ribosomal, mitochondrial, and non-variable genes (21). CRISPRclean lowered the proportion of ribosomal and mitochondrial genes in the PBMC library by over 30% (Figure 6A).

**Fig. 6.**
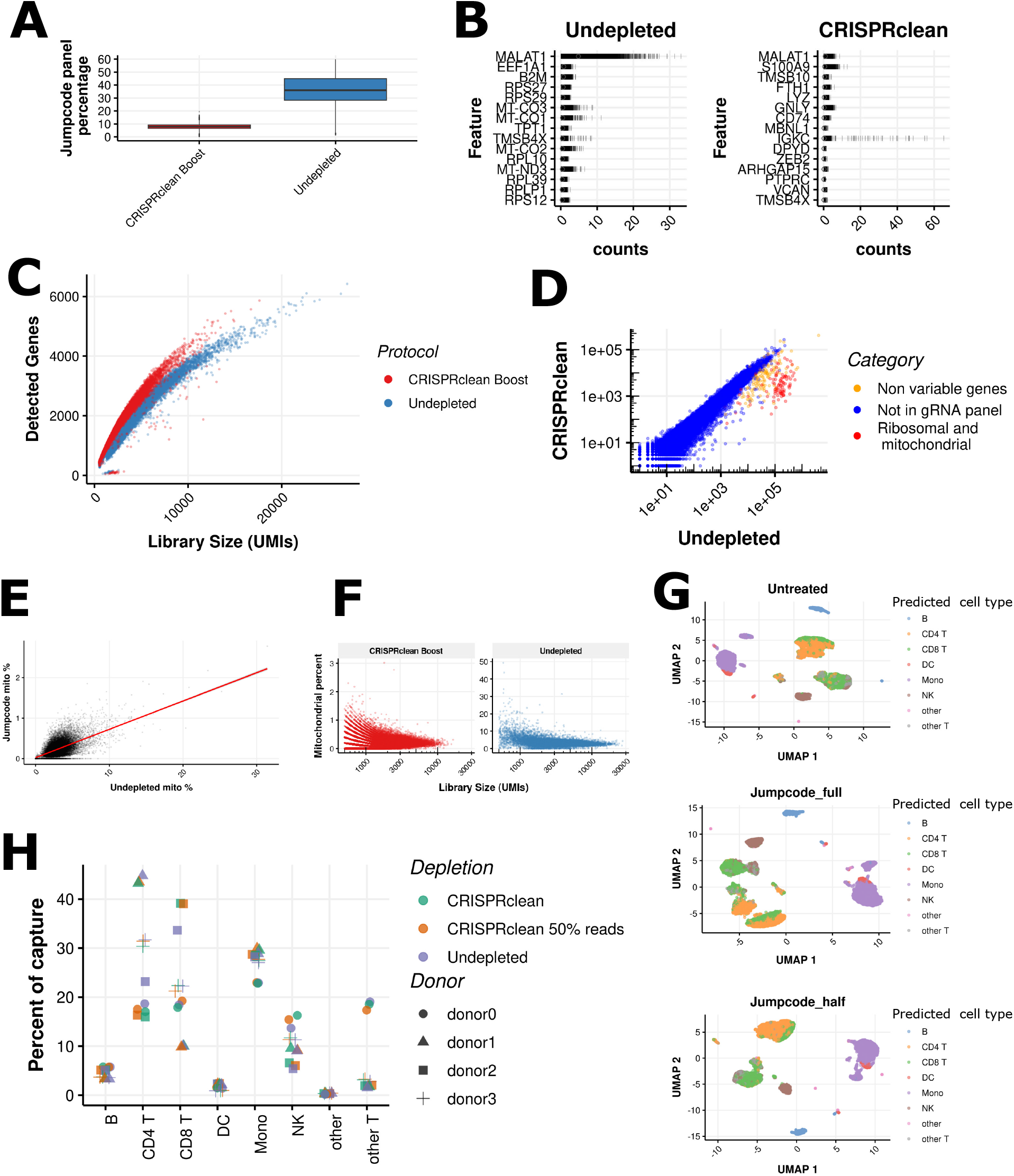
Evaluation of Jumpcode CRISPRclean Single Cell RNA Boost Kit in MULTI-Seq LMO PBMCs. (A) Percentage of counts on genes in the CRISPRclean panel. The gene expression library was sequenced with and without CRISPRclean treatment. (B) Top 20 highly expressed genes in CRISPRclean treated and untreated libraries. Genes prefixed as “RP” or “MT-” represent ribosomal or mitochondrial genes, respectively. (C) Relationship between library size and number of detected genes per cell. (D) Gene expression comparison, each dot represents the sum of counts across all single cells for a gene. (E) Correlation of mitochondrial gene percentages. The trend line is a linear fit. (F) Relationship between library size and mitochondrial gene percentage. (G) UMAP dimension reduction based on gene expression data. The CRISPRclean library was additionally downsampled to 50% of the untreated library. (H) Summary of immune subsets recovered from untreated, CRISPRclean treated, and downsampled CRISPRclean treated gene expression data.

Following CRISPRclean treatment, ribosomal and mitochondrial genes were eliminated from the list of most highly expressed genes, and *MALAT1* was considerably depleted (Figure 6B). The enhancement in gene detection was moderate (Figure 6C), while off-target depletion of genes remained minimal (Figure 6D). Interestingly, unlisted ribosomal genes in the CRISPRclean panel were also degraded, potentially due to homology.

The per-cell percentage of counts for mitochondrial genes is a common quality control metric in scRNA-Seq analysis (22). We investigated whether this metric remains reliable for removing low-quality cells in CRISPRclean-depleted samples (Figure 6E and F). Although a positive correlation between untreated and depleted mitochondrial gene percentages was observed, a distinct subset of cells was removed based on data-driven outlier-based quality control. Apart from mitochondrial gene expression, no apparent patterns in the expression of other genes distinguished untreated and depleted cells.

We next confirmed that cell type recovery remained unaffected by the depletion of ribosomal and mitochondrial genes (Figure 6G). We also examined whether cluster resolution improved or if the ability to annotate immune subsets was enhanced. Following clustering and annotation, no difference in the composition of CRISPRclean-depleted libraries was observed (Figure 6H).

## Discussion

Our study compared various sample multiplexing reagents for scRNA-Seq experiments. We limited our comparison to commercially available technologies that do not require genetic manipulation of cells. For PBMCs, we found MULTISeq LMO to be the superior reagent. Hashtag antibodies also performed well, with the exception of a single tag oligo HTO_1 in this particular experiment. Potential explanations include the formation of antibody aggregates, which is mentioned on the frequently asked questions page of the 10x Genomics website. Reduced signal-to-noise ratios might also be attributed to incomplete removal of supernatants during wash steps.

CellPlex at the manufacturer’s recommended concentration, exhibited the poorest performance due to high background.

By titrating and diluting the CellPlex reagent ten-fold, the signal-to-noise ratio was improved. The additional background was likely introduced during storage of the pooled sample prior to single-cell capture, as the concentration of the CellPlex reagent is higher than for MULTI-Seq or hashtag antibody. Any passive transfer of multiplexing oligo tags by diffusion would be more pronounced for a concentrated reagent. The mechanism of tag oligo exchange between cells or between nuclei remains unclear. In our observations, it occurred in both high-quality cell lines and lower-quality dissociated tissues in a time dependent fashion. Based on 10x Genomics guidelines to maintain samples on ice after pooling, passive diffusion is the likely cause. This phenomenon could be further investigated using live cell fluorescent microscopy.

We caution against the use of sample multiplexing reagents particularly for fragile samples. labeling of cells with multiplexing oligos necessitates additional sample incubation and washing steps. These additional manipulations can compromise the viability of fragile cell types such as mouse embryonic brain and tumour nuclei. The benefits of sample multiplexing with respect to cost and batch effect minimization should be assessed against the risk of reductions in data quality. Large-scale experiments involving upwards of 96 samples multiplexed together have only been demonstrated using cell lines (7); However, our results suggest that it would be challenging to achieve such a scale with primary cells or tissues.

Comparing the assignments based on sample multiplexing oligos with SNP assignments confirmed the accuracy of all labeling strategies. Fewer than 2% of cells were misclassified to the wrong donor in the PBMC experiment. Nevertheless, SNP assignments operated with higher cell recovery with fewer cells being lost to an unassigned category. We recommend the use of SNPs over sample multiplexing reagents where applicable. In our algorithmic comparison, we found that all methods performed similarly when data quality is high as was the case in the PBMC experiment. This is consistent with other studies (23). Where the algorithms diverge in terms of performance is on lower quality data. Here we found BFF_Cluster could be used to rescue more cells, at the expense of an elevated false positive rate. Combining demultiplexing algorithms with orthogonal information, such as SNP genotypes or endogenous gene expression, may be a useful strategy to rescue poorly performing datasets, particularly with fragile samples where signal-to-noise is sub-optimal. It is important to note that we only tested algorithms available within the cellhashR package (14). As new algorithms are constantly emerging (24, 25), dedicated benchmarking studies focusing exclusively on bioinformatics analysis will be necessary (23). Our primary focus was on laboratory elements that can enhance demultiplexing performance.

An advantage of the fixed scRNA-Seq kits from 10x Genomics and Parse Biosciences is that sample multiplexing is embedded in the molecular biology, with no additional sample handling required as for sample multiplexing oligos. While the Parse Biosciences kit has a lower doublet rate, the doublets in the 10x Genomics product are usable. For large experiments where gene expression is the sole readout, we expect that fixed RNA kits will become dominant. For PDX nuclei, we found that the Parse Biosciences kit was more reliable, whereas 10x Genomics Flex suffered from recurrent wetting failures. The greater propensity of nuclei for aggregation and clumping compared to intact cells may explain these failures encountered by the microfluidics-reliant 10x Genomics technology. It is also possible that the higher nuclei input may have also contributed to the formation of wetting failures. In the multiplexing configuration of 10x Genomics Flex, droplets containing heterotypic doublets are usable, therefore we input a higher number of nuclei into the microfluidics device compared to conventional 3’ chemistry. This overloading feature is built into the pricing structure, where Flex is more expensive than 5’ and 3’ kits, if overloading is not utilized.

CRISPRclean offers a promising approach to focusing sequencing resources on specific genes of interest. In our assessment of PBMCs, which represent the sample type used in the manufacturer’s demonstration, we found that the technology is consistent with the manufacturer’s claims. However, when applied to PDX samples, we observed a less pronounced re-focusing of sequencing effort. This discrepancy can be ascribed to the diminished expression of genes contained within the gRNA panel in PDX samples compared to PBMCs. While we conducted a comparison of cell type recovery, we did not perform an in-depth analysis on the impact of normalization, which typically relies on non-variable genes. In our data, we identified off-target depletion of genes, some of which have been previously reported (21). The remaining off-target effects could be attributed to statistical noise and additional replicate experiments are necessary to evaluate their reproducibility. Following our assessment of CRISPRclean, a new version of gRNAs has been introduced, addressing known off-target effects and potentially enhancing the specificity of this technology.

In conclusion, our findings indicate that the choice and extent of multiplexing for scRNA-Seq should be contingent on the type of sample under investigation. Samples composed of cells that withstand *ex vivo* manipulation can accommodate a high degree of multiplexing. Conversely for delicate samples, it is more prudent to minimize multiplexing and instead invest in additional consumable costs. This approach ensures the preservation of high data quality.

## Methods

All oligonucleotides were purchased from Integrated DNA Technologies. Sequences are provided in Table S5.

### Ethical statement

PBMCs were isolated from unrelated healthy control donor samples obtained from the Volunteer Blood Donor Registry (VBDR, WEHI). Informed consent was obtained from all individual participants prior to inclusion in the study. The study was performed according to the principles of the 1964 Helsinki declaration and its later amendments and was approved by local Human Research Ethics Committee (WEHI Approved project 10/02). All experiments involving animals were performed according to the animal ethics guidelines and were approved by the WEHI Animal Ethics Committee (2019.024).

### PBMC sample multiplexing labeling

#### CellPlex

Labeling was performed according to 10x Genomics demonstrated protocol CG000391 Rev A “Cell Multiplexing Oligo Labeling for Single Cell RNA Sequencing Protocol” with an input of 250,000 cells. Library preparation was performed according to CG000388 Rev A “Chromium Next GEM Single Cell 3’ Reagent Kits v3.1 (Dual Index) with Feature Barcode technology for Cell Multiplexing”.

#### MULTI-Seq

The MULTI-Seq protocol (7) was followed with 200nM of each anchor-barcode complex being used in the labeling step. The poly-A capture sequence was replaced with the 10x Genomics feature barcode capture 2 sequence. This required the library preparation PCR to be performed with the Nextera read 1 primer instead of the TruSeq read 1 primer. PCR conditions otherwise remained the same.

#### Total-Seq A Hashtag antibody

The Biolegend protocol for Total-Seq A hashtag labeling was followed with the exception that ten-fold less antibody was used, 0.1 *μ*g per labeling reaction. Each multiplexing protocol was captured on a separate 10x Genomics lane to avoid ambient oligo effects.

### Mouse embryonic brain sample preparation

RosaERT2Cre/RosaERT2Cre mice were intercrossed with Loxcode/Loxcode mice (26, 27). Presence of a vaginal plug was used to determine day of conception. Pregnant dam was induced with 50*μ*g of 4 hydroxytamoxifen injected intravenously at day 7.5 of pregnancy. Embryos were collected at E18.5. After decapitation, the brain was dissected and processed for enzymatic dissociation. Embryos were dissected with the 10x Genomics protocol CG00055 Rev C “Dissociation of Mouse Embryonic Neural Tissue for Single Cell RNA Sequencing” with the following modifications: Embryos were dissected in Hibernate-E Medium media (ThermoFisher A1247601), supplemented with 1% B27 (ThermoFisher 17504044) and 1x GlutaMAX (ThermoFisher 35050061). Benzonase Nuclease (Millipore E1014) was added to Papain (Millipore P4762) at a dilution of 1:5000 during dissociation.

### Mouse E18.5 brain sample multiplexing experiment

100,000 cells were aliquotted per well across 12 wells of a round bottom 96 well plate (Falcon 353077). To label cells at high throughput, a combination of 10x Genomics protocols CG000391 Rev B “Cell Multiplexing Oligo Labeling for Single Cell RNA Sequencing Protocols with Feature Barcode technology” and CG000426 Rev A “High Throughput Sample Preparation for Single Cell RNA Sequencing” were utilized.

#### CellPlex

CellPlex oligos were diluted 1:10 in PBS prior to incubation with 100,000 cells at room temperature. After 5 min, 200 *μ*L of PBS + 1% BSA was added and samples centrifuged at 300g for 4min at 4 °C. The supernatant was aspirated with a multichannel pipette, leaving approximately 10 *μ*L of supernatant. A single wash of 200 *μ*L PBS + 1% BSA was performed prior to pooling and sorting by flow cytometry. Library preparation followed the 10x Genomics CG000388 Rev A protocol with no alternation of PCR cycle number.

#### MULTI-Seq lipid modified oligo (LMO)

labeling was performed at half volume (100 *μ*L total) to avoid overflow of the round bottom 96 well plate. 200nM of anchor-barcode complex was used in the labeling step with 100,000 cells for 5 minutes on ice. Co-anchor oligo was then added for a further 5 minutes on ice prior to quenching with 200 *μ*L PBS. A single wash was performed prior to flow cytometry. Library preparation followed the same workflow as for the PBMC experiment.

#### Custom MULTI-Seq cholesterol modified oligo (CMO)

The MULTI-Seq LMO process was followed, substituting anchor and co-anchor oligos with a custom cholesterol modified oligo (CMO) containing the Nextera read 2 PCR handle (Supplementary table 5). Since the oligo was designed for compatibility with the CellPlex workflow, library preparation followed the 10x Genomics CG000388 Rev A protocol without PCR cycle alterations.

#### 10x Genomics capture

After labeling with multiplexing tag oligos, individual samples were pooled at equal volumes without a cell count. The pooled single-cell suspension was counted and diluted to a final concentration of 812 cells per *μ*L, aiming to load 35,000 cells into each lane and obtain 20,000 barcode-containing droplets at a 16% theoretical doublet rate. 10x Genomics v3.1 dual index kits were used. Each multiplexing protocol was captured on a separate 10x Genomics lane to avoid ambient oligo effects.

### Ovarian carcinosarcoma xenograft fresh nuclei experiments

PDX models were established through transplanting fragments of tumor tissue obtained from patients consented to the WEHI Stafford Fox Rare Cancer Program (28). Following ethical endpoint and tumor dissection, rice-sized tissue pieces were snap-frozen on dry ice and stored at -80°C until processing. Single nucleus suspensions were generated from frozen tissue pieces using the Chromium Nuclei Isolation Kit with RNase inhibitor (PN-1000494), following user guide CG000505 Rev A. As input tissue pieces weighed over 50mg, the four tumor pieces from each donor were cut in half, and the nucleus preparation was performed in duplicate. *10x Genomics unlabelled capture*. To examine the effects of extended storage time on nuclei integrity a capture was performed prior to any multiplexing labeling step, approximately 90 minutes before the labelled samples were captured. The single nuclei suspensions from each PDX donor were counted and pooled to a final concentration of 692 nuclei per *μ*L to load 30,000 nuclei and obtain 17,177 barcode containing droplets at 13.82% theoretical doublet rate.

#### CellPlex

CellPlex oligos were diluted 1:10 in PBS prior to incubation with 250,000 cells at room temperature. After 5 minutes of labeling time at room temperature, 200 *μ*L of PBS + 1% BSA was added and samples centrifuged at 300 g for 4min at 4 °C. The supernatant was aspirated with a multichannel pipette, leaving approximately 10 *μ*L of supernatant. A single wash of 200uL PBS + 1% BSA was performed prior to pooling and sorting by flow cytometry. Library preparation followed the 10x Genomics CG000388 Rev A protocol with no alternation of PCR cycle number.

#### Custom MULTI-Seq custom cholesterol modified oligo (CMO)

The same process for MULTI-Seq LMO was performed, except for the substitution of the anchor and co-anchor oligos for a custom cholesterol modified oligo (CMO) bearing the Nextera read 2 PCR handle. As the oligo was designed to be compatible with the CellPlex workflow, library preparation followed the 10x Genomics CG000388 Rev A protocol with no alteration of PCR cycles.

#### TotalSeq A anti-Nuclear Pore Complex Antibody

The Biolegend Protocol for Total-Seq A hashtags was followed with 1 *μ*g of antibody per labeling reaction (accessed 22 August 2022).

### Ovarian carcinosarcoma xenograft fixed RNA experiments

To obtain sufficient nuclei to run the same suspension across both fixed kits we used EZ lysis buffer (Sigma NUC101), at the expense of greater debris. 500 *μ*L of lysis buffer was added to approximately 50*μ*g frozen tissue pieces. Sample was titruated with wide bore p1000 tips until homogenized. Nuclei were incubated on ice for 5 minutes followed by centrifugation at 500 g for 5min at 4°C. The supernatant was removed followed by two washes in 1 mL PBS + 1% BSA. Nuclei samples were then divided in half and fixed with manufacturer specific protocols and reagents.

#### Parse Biosciences Evercode version 2 on PDX nuclei

Between 150,000 and 500,000 nuclei were fixed and stored at -80°C for 6 weeks prior to processing using the Parse Biosciences Evercode WT Mini v2 kit (ECW02010) version 2.0.0 protocol. After visual inspection of the nuclei following fixation, 2 of the 4 samples exceeded the maximum clumping parameters and were omitted, leaving 2 remaining samples. Each sample was then processed in 2 wells of a version 2 mini kit following the manufacturer’s guidelines. Two sublibraries of 5,000 cells underwent the downstream cell lysis and PCR amplification steps.

### Jumpcode CRISPRclean depletion

The MULTI-Seq LMO library from the PBMC experiment and unlabelled library from the ovarian PDX experiment were treated with Jumpcode CRISPRclean Single Cell RNA Boost Kit, (KIT1018) according to the manufacturer’s instructions.

### Bioinformatics analysis

All downstream analysis was performed in R version 4.2.1 (29). The code, data and analyses used to generate these figures is available from GitHub. Each multiplexing labeling protocol evaluated was treated as a separate dataset without integration. Tabular data was manipulated with the tidyverse package (30).

Cell annotation was performed with Seurat version 4.0.6, TransferData function (31). The reference for human PBMCs was provided with Seurat multimodal reference mapping vignette. The reference for mouse E18.5 brain was (32), subsetting cells for the E18.5 timepoint.

Doublet detection based on gene expression was performed with demuxafy version 1.0.3 (16). The majority vote from the output of DoubletFinder, scDblFinder, scds and scrublet was used to assign multiplets and singlets. Ambient RNA estimation was performed with Souporcell version 2.0 (33).

## Statistical analysis

Differential cell type abundance analysis was performed by summarising the number of cells la-belled with a given cell type annotation for each sample of origin, followed by testing for differences in the abundance of cell types between cell demultiplexing protocols with the edgeR package (34).

For differential gene expression analysis single-cells were first aggregated to pseudobulks based on CRISPRclean treatment with the aggregateAcrossCells function from the scuttle package (35). The edgeR package was then used to compute differentially expressed genes.

## Data statement

The count matrices and metadata are available as SingleCellExperiment objects at Zenodo, DOI: 10.5281/zenodo.8031078

## Supplementary Information

Supplementary information includes additional results, methods and discussion. Supplementary tables 1 - 5 and supplementary figures 1 - 10.

## ACKNOWLEDGEMENTS

We thank all those individuals that donated the samples that enabled this study. We acknowledge the WEHI Genomics, Flow Cytometry Facility and Animal Bioservices for professional and timely service.

D.V.B is supported by funding to the Advanced Genomics Facility from the Walter and Eliza Hall Institute. S.A.B. is supported by a Victorian Cancer Agency MidCareer Research Fellowship (MCRF22003), Z.M. is supported by the Greg Lange Memorial Postdoctoral Fellowship, S.F. is supported by a National Health and Medical Research Council of Australia (NHMRC) Ideas Grant (1184421). V.L.B is supported by Sir Clive McPherson Family Fellowship and D.W Keir Fellowship. This work was financially supported in part through the authors’ membership of the Brain Cancer Centre, support from Carrie’s Beanies 4 Brain Cancer, a Priority-Driven Collaborative Cancer Research Scheme Grant funded by Cancer Australia to S.A.B. (2003127).

Experimental design figures were created using Biorender.com. This preprint is formatted using a LATEXclass by HenriquesLab.

## AUTHOR CONTRIBUTIONS

D.V.B, P.F.H,D.A.Z,R.B conceptualized the study and designed experiments.

D.V.B,C.A,R.M,C.B,A.H performed sample preparation.

D.V.B,C.A,L.L,T.M.B,S.W.Z.O,Z.M generated single cell libraries.

D.V.B,P.G performed bioinformatics, data processing, and computational analyses based on code from P.F.H and A.D.

S.M,and S.W generated sequence data from supplied libraries.

Critical reagents and resources were provided by A.J.F,V.L.B,S.T,S.H.N,T.S.W,G.D,H.E.B,C.L.B,T.P,C.L.S,S.A.B,J.R.W,S.A.B.

D.V.B wrote the manuscript with approval from all of the authors.

## Declaration of generative AI use in scientific writing

The initial rough draft of introduction, results and discussion sections were prepared by D.V.B without AI assistance. ChatGPT Plus (GPT-4) was subsequently used for copy editing of each paragraph with the prompt “*act as a scientific copy editor and suggest improvements to my manuscript. List each change that you suggest*.” The edits were manually assessed before accepting or rejecting. ChatGPT Plus was then used to generate a 200 word abstract based on the results and discussion section. The output was manually reviewed and edited by D.V.B before inclusion and take they full responsibility for the content in the manuscript. Finally, ChatGPT was asked to provide five manuscript titles and keywords based on the results and discussion sections.

## Supplementary Results

### Immune lineage and subset analysis

After annotating the singlet-containing droplets, we noticed a difference in oligo tag library size between lymphoid and myleoid immune subsets especially for CellPlex (Figure S4E). This difference was not statistically significant and there was no difference in immune subset recovery across broad (Figure 2F) and finer (Figure S4F) cell annotations. This effect is likely to be due to differences in cell size between lymphoid and myleoid cells.

### Comparison of diluted and undiluted Cellplex in mouse embryonic brain experiment

We adopted the same experimental setup as the Cellplex Demonstration Data (v3.1 Chemistry) from 10x Genomics to facilitate a comparison with undiluted CellPlex (Figure S**??**B). The signal-to-noise ratio remained comparable, although a slightly superior assignment of cell-containing droplets to tags was observed in the undiluted CellPlex dataset (Figure S5C). As these two datasets were generated using different samples and in distinct laboratories, it is challenging to ascertain whether this effect is attributable to reagent dilution. Notably, given the significantly lower background in the diluted CellPlex library, fewer resources would be expended on sequencing through background multiplexing oligos.

### 10x Genomics Flex on intact Ovarian PDX single-cell suspension

Following the experiment conducted on PDX tumor nuclei, we carried out an additional 10x Genomics Flex experiment utilizing intact cells. We hypothesized that nuclear leakage contributed to cell clumping, and that this issue could be mitigated by working with cells. Two methods of preparing single-cell suspensions were evaluated: firstly, fixing a freshly prepared single-cell suspension created through enzymatic tissue dissociation, and secondly, fixing the tissue prior to dissociation.

While both methods successfully yielded single-cell suspensions with minimal clumps, microfluidic clogs occurred in two independent captures, resulting in a final Cell Ranger multi estimate of 1,200 cells rather than the theoretical 80,000. The quality of data acquired from tissue fixation prior to dissociation was significantly inferior to that obtained by fixing a freshly generated cell suspension (Figure S8L), exhibiting poor gene expression correlation (Figure S8N).

### Further CRISPRclean evaluation in PBMCs

As reported in Jumpcode advertising materials, CRISPRclean treatment may enhance the detection of additional sub-populations that could be otherwise obscured by the presence of nonvariable genes. Contrary to the manufacturer’s claims, we observed no difference in the abundance of classical and nonclassical monocytes (Figure S9A). In both libraries, a clear distinction was noted between cells expressing *FCGR3A* and *CD14* (Figure S9C). We compared gene expression between CRISPRclean-treated and untreated samples and identified fewer than 50 differentially expressed genes (Figure S9B). A subset of these genes overlapped with known off-target genes, as reported by Pandey et al., which are attributed to gRNA homology (21).

### CRISPRclean evaluation in Ovarian PDX

We also evaluated the CRISPRclean kit in ovarian carcinosarcoma PDX nuclei, where its effects were diminished, likely due to the lower baseline expression of ribosomal and mitochondrial genes in this sample (Figure S10A). We analysed differential gene expression between CRISPRclean-treated and untreated samples in both experiments and identified fewer than 60 (Figure S10F). Approximately half of these overlapped with known off-target genes reported in Pandey et al., resulting from gRNA homology (21).

## Supplementary Discussion

### Choice of high throughput labeling protocol for mouse embryonic brain experiment

The decision to sort the mouse embryo experiment post-labeling was influenced by the guidelines provided by 10x Genomics and a direct comparative study of preand post-label sorting by Mylka et al. (13). Given the contrived nature of our experiment, which entailed a comparison of different multiplexing reagents on the same sample within a single day, sequential FACS sorts for varying protocols were necessary. This arrangement, however, resulted in MULTI-Seq samples being kept on ice for a longer duration than CellPlex.

In a more typical experimental setup, only a single sorting step would be required post-pooling of samples. Our findings suggest that swift cell capture is crucial to prevent loss of viability and to minimize multiplexing oligo tag exchange. We recommend having all the necessary reagents for 10x Genomics captures prepared during the sorting process, thus enabling single-cell partitioning to occur as promptly as possible.

### Cell demultiplexing algorithm comparisons

BFF_Cluster demonstrated variable performance across the different datasets in our study. This tool utilizes a non-parametric, quantile-based normalization procedure, which assumes a bimodal distribution in the oligo tag count data (14). The primary goal is to enhance the signal-to-noise ratio in under-performing datasets. Notably, in the CellPlex PBMC dataset and mouse embryo datasets, BFF_Cluster identified the highest number of cells. However, in the PBMC dataset, where we had SNP ground truth as reference, BFF_Cluster produced the most false positives (Figure 2E). The majority of these incorrect assignments were multipletto-singlet errors. These could potentially be mitigated through additional filtering using gene expression-based doublet detection software like Demuxafy. We implemented this strategy in the mouse embryonic brain experiment where the signal-to-noise ratio was low, yet some signal was detectable. The aim of this experiment was to overload the 10x Genomics capture process to maximize the number of cells captured. In contrast, in the Ovarian PDX nuclei experiment, BFF_Cluster did not yield any droplet calls for either the Hashtag antibody or MULTI-Seq CMO. This conservative behaviour is more desirable in instances of experimental failure.

In the Ovarian PDX experiment, Seurat HTO_Demux() yielded the highest number of cells, with an even distribution of cells recovered per multiplexing tag. This might be attributable to the k-medoids mediated clustering step that explicitly models 1 plus the number of tags used (23). Conversely, the alternative demultiplexing algorithms, which did not implicitly account for the number of tags used in the experiment, were more conservative, labeling the majority of nuclei as unassigned. It is worth noting that HTO_Demux() is a heuristic algorithm and its sensitivity is subject to the user-defined quantile threshold, which in our study was left at the default value of 0.99 (36).

### Fixed snRNA-Seq experiments

Our experiments suggest Parse Biosciences Evercode is particularly well suited for fragile samples, with a higher number of reads in cells and molecules and genes detected overall. The multiple washing steps between split pool rounds likely remove ambient RNA plus the split pool strategy also makes it unlikely that ambient RNA will follow the same barcoding path as intact nuclei.The lack of microfluidics may also reduce shear stress on nuclei during sample handling.

The disadvantage of Parse Biosciences is a more labour intensive workflow than 10x Genomics Flex with an entire day in the laboratory to generate barcoded single-cells compared to less than half this time for 10x Genomics Flex in our experience.Handling multichannel pipettes across many microwell plates may also contribute to high inter-operator variability compared to an automated microfluidic instrument. Currently both multiplexed Fixed RNA kit are not compatible with antibodies and Flex is incompatible with CRISPR experiments. Where these modalities are required, sample multiplexing oligos will continue to be useful.Through a splint oligo or gap filling reaction we anticipate that these additional modalities will be enabled in the future.

### CRISPRclean experiments

It is crucial to consider that CRISPRclean treatment adds time and expense to library preparation and sequencing. In our PBMC experiment we did not observe an obvious benefit in cluster identification, though the claimed cost reduction was achieved. While the PBMC sample benefited from CRISPRclean depletion, in the PDX sample, the cost of the CRISPRclean treatment outweighed the reduction in sequencing cost (approx $350 cost vs $210 AUD saved).We anticipate that for samples where non-variable genes make up a high proportion of the sequencing library, CRISPRclean will be integrated into a routine laboratory workflow.

The choice of scRNA-Seq chemistry should also be factored in when deciding to use CRISPRclean. We observed a lower proportion of ribosomal and mitochondrial genes in the Parse Biosciences library compared to the 10x Genomics v3.1 library and would not have recovered the cost of CRISPRClean treatment in the Parse Biosciences experiment.

## Supplementary Methods

### Titration of multiplexing reagents by flow cytometry

K562 cells were used for flow cytometry experiments. Single-cell suspensions were labeled with the relevant multiplexing reagents, with the addition of 200nM of fluorescent detection oligo during the labeling step. These detection oligos have the same sequence present on the 10x Genomics 3’ v3.1 gel bead, comprising either the poly-A or feature barcode 2 capture sequence with a 5’ Alexa 647 modification (Table S5). Following washing steps, the singlecell suspensions were resuspended in 500 *μ*L PBS 1% BSA, supplemented with 0.1*μ*g/mL 4’,6-diamidino-2-phenylindole (DAPI, ThermoFisher 62248) as a viability dye. Cells were analyzed on a BD LSR II flow cytometer. Data analysis was performed using the ggCyto package in R (37).

### PBMC isolation and cryopreservation

PBMCs were isolated from fresh whole blood by density gradient Ficoll-Leucosep centrifugation. Cells were cryopreserved in liquid nitrogen in FCS containing 10% DMSO and placed in -80°C freezer for a short-term storage. Vials were then transferred to liquid nitrogen vapour phase tanks for long-term storage (38).

### PBMC thawing sample preparation

Cryopreserved PBMC samples were rapidly thawed in a 37°C water bath. Each sample was transferred into a 15-ml tube using a p1000 *μ*l tip without mixing by pipetting. Next, 1 ml of 37°C pre-warmed media (RPMI supplemented with 10% FCS) was added dropwise with gentle swirling of the sample. After 1 min, an equal volume of media was added; this process of adding an equal volume of media was repeated a further 3 times. The samples were then centrifuged at 400g for 5 min at 4°C. Pellets were resuspended in 1mL of RPMI supplemented with 10% FCS, and filtered with a 40 *μ*M strainer (Falcon Cat 352340). DAPI was added to a final concentration of 0.1 *μ*g/mL prior to the sorting of viable (DAPI negative) cells. Cells were collected in cell staining buffer (Biolegend 420201) and counted with Countess III FL (ThermoFisher AMQAX1000). Each of the PBMC donor samples was split into aliquots of 250,000 cells each prior to sample multiplexing oligo labeling.

### Mouse E18.5 brain sample multiplexing experiment

To compare CellPlex reagent diluted ten-fold versus undiluted, Cellplex Demonstration Data (v3.1 Chemistry) *“30k Mouse E18 Combined Cortex, Hippocampus and Subventricular Zone Cells Multiplexed, 12 CMOs”* was downloaded from the 10x Genomics website.

### Ovarian carcinosarcoma xenograft fixed RNA experiments

#### 10x Genomics Flex on PDX nuclei

The same nucleus suspension used for Parse Biosciences was fixed with 10x Genomics Flex-specific fixative. Between 150,000 to 500,000 nuclei were fixed and stored at -80°C for 2 weeks prior to processing with 10x Genomics Flex kit, CG000527 Rev B. Initially, the nuclei were observed to be in a single suspension. However, following probe hybridisation and multiple wash steps, the clumping rate increased. The sample was filtered twice with a 30*μ*m pre-separation filter (130-041-407, Miltenyi Biotech) prior to the final cell count. After capture with the Chromium X controller, the volume of GEMs was lower than expected, indicating a wetting failure. The library was sequenced, yielding 30M reads and a dataset of 303 cells with a reads-in-cells metric of 29%. Due to the low cell yield, the dataset was not analysed further.

#### 10x Genomics Flex on PDX intact cells and tissue

To obtain fixed cells, tumour pieces were dissociated with collagenase, dispase, and DNase, and two million cells were fixed with the 10x Genomics-specific fixative. At the same time, approximately 50 mg of rice-sized tissue pieces were fixed directly and stored at 4°C for 5 days according to the manufacturer’s instructions. Fixed tumour pieces were then thawed and dissociated with collagenase, dispase, and DNase. Single cells were immediately processed, along with cells that had been dissociated and then fixed with the 10x Genomics Flex kit CG000527 Rev B. A minor wetting failure was noted, with approximately 90 *μ*L recovery of GEMs (80% of expected volume). The remaining cell suspension was used for a second Chromium X run performed with another microfluidics chip, yielding the same outcome.

### Sequencing

Sequencing for PBMC samples was performed on a NextSeq500 targeting 5,000 reads per cell for the multiplexing oligos and 20,000 reads per cell for the gene expression. An initial sequencing run of 28 bp (read 1), 10 bp (index read 1), 10bp (index read 2), 90b p (read 2) was performed, followed by a second run of 28 bp,10 bp,10 bp,44 bp for the multiplexing oligos only. A further aliquot of PBMC gene expression libraries were sequenced with MGI DNBSEQ-G400 instrument. Standard 10x Genomics libraries were converted with MGIEasy Universal Library Conversion kit (App-A, 1000004155). 25 ng of standard gene expression library was used as input to the conversion step. 8 PCR cycles were performed prior to circulisation. Sequencing was performed with the DNBSEQ-G400RS High-Throughput Sequencing Kit: FCL PE100, (1000016949), with MGISEQ-2000RS Sequencing Flow Cell v3, (000008403).

Mouse E18.5 brain samples were sequenced on a NextSeq2000 P3 flow cell targeting 2,000 reads per cell for the multiplexing oligos and 16,500 reads per cell for the gene expression. The ovarian carcinosarcoma nuclei 10x Genomics v3.1 experiment was sequenced on a NextSeq2000 P3 flow cell. For the unlabelled sample a mean of 20,000 reads per nucleus were targeted (12,000 reads in cells). For the samples labelled with multiplexing oligos 3,500 reads per cell for oligos and 5,000 reads per cell for the gene expression were targeted. Given the poor performance of the labelled nuclei further sequencing was not performed. The ovarian carcinosarcoma fixed nuclei Parse Bioscience v2 experiment was sequenced on a MGI DNBSEQ-G400 instrument following library conversion with App-A.300,000 reads per cell were targeted. Reads were then downsampled to a comparable level to the 10x Genomics unlabelled sample (16,500 reads per cell). The ovarian carcinosarcoma 10x Genomics Flex v1 experiment was sequenced on a NextSeq2000 P2 flow cell. We targeted 40,000 cells at 10,000 reads per cell. Given the sample clog, only 12,000 cells were recovered, at 430,000 reads per cell.

### Bioinformatics preprocessing

#### PBMC and mouse embryo experiment

Sequencing data was demultiplexed with Cell Ranger v6.0.0 mkfastq. MGI sequencing data was demultiplexed with splitBarcode v2.0.0 from MGI. Production of count matrices and demultiplexing of samples was performed with Cell Ranger multi v6.0.0 using 10x Genomics pre-built GRCh38 reference genome and transcriptome for PBMCs and mm10 for mouse embryo (2020-A July 7, 2020 version). Demultiplexing of PBMC samples by genotype was performed with cellsnp-lite v1.2.0 (39), with 36.6M SNPs with minor allele frequency (MAF) > 0.0005 in the 1000 Genomes Project. SNP vcf file. Then vireo was used to assign each cell barcode to 1 of 8 donors, doublets, or unassigned based on SNP genotypes (40). This was performed separately for each multiplexing protocol followed by global donor matching in aggregate based on the genotype profile of each donor.

#### Ovarian carcinosarcoma nuclei 10x Genomics v3.1

Sequencing data was demultiplexed with Cell Ranger v7.0.0 mkfastq. Production of count matrices and demultiplexing of samples was performed with Cell Ranger multi v 7.0.0 using 10x Genomics pre-built GRCh38 and mm10-2020-A reference genome and transcriptome (2020-A (July 7, 2020)). Demultiplexing of samples was performed with cellsnp-lite and vireo as for the PBMC dataset.

#### Ovarian carcinosarcoma nuclei Parse Biosciences mini v2

Data was processed with Parse Biosciences spilt-pipe v1.0.3 using a mixed GRCh38 and mm10 reference genome and transcriptome with default parameters.

#### Ovarian carcinosarcoma nuclei 10x Genomics Flex

Data was processed with Cell Ranger multi v7.0.0.

## Supplementary Tables

**Supplementary Table 1.** Summary of sample multiplexing reagents

| Name | Composition | Compatibility | Plexing | Cost per sample |
| --- | --- | --- | --- | --- |
| Total-Seq hashtag | Antibody | Cells OR nuclei | 15 | \$\$ |
| MULTI-Seq | Lipid | Cells OR nuclei | Unlimited | \$ |
| CellPlex | Lipid | Cells AND nuclei | 12 | \$\$\$ |

**Supplementary Table 2.** Metrics from the mouse E18.5 brain multiplexing experiment.

|  | CellPlex | MULTI-Seq CMO | MULTI-Seq LMO |
| --- | --- | --- | --- |
| Cell containing droplets | 27,125 | 19,494 | 9,745 |
| Theoretical Doublets | 5,886 | 3,040 | 759 |
| Theoretical doublets % | 21.70% | 15.60% | 7.80% |

**Supplementary Table 3.** Metrics from the fresh Ovarian carcinosarcoma PDX experiment.

|  | CellPlex | Hashtag Ab | MULTI-seq CMO |
| --- | --- | --- | --- |
| Estimated Number of Cells | 5,509 | 7,971 | 5,156 |
| Mean Reads per Cell | 5,746 | 7,320 | 11,235 |
| Median UMI Counts per Cell | 157 | 352 | 650 |
| Sequencing Saturation | 48.40% | 52.40% | 50.30% |
| Median Genes per Cell | 148 | 321 | 558 |
| Reads Mapped Confidently to Transcriptome | 34.30% | 49.80% | 42.70% |

**Supplementary Table 4.** Metrics from the fixed Ovarian carcinosarcoma PDX experiment.

|  | Parse v2 mini | 10x Genomics v3.1 |
| --- | --- | --- |
| Cells recovered | 3,683 (1/3 kit) | 16,181 (1 capture) |
| Doublet rate (Heterotypic) | 0.73% | 7% |
| Reads per cell | 16,587 | 12,817 |
| Sequencing saturation | 5.5% | 54.3% |
| Reads in cells | 84.2% | 50.8% |
| Transcriptome mapping | 72.8% | 47.3% |

Supplementary Table 5. which contains oligonucleotide sequences is provided as an Excel file.

## Supplementary Figures

**Supp figure 1**. Optimisation of the sample multiplexing tag oligos by flow cytometry.

**Supp figure 2**. Related to Figure 1. Additional metrics on the performance of multiplexing tag oligo reagents in PBMC experiment.

**Supp figure 3**. Related to Figure 1. Titration of CellPlex lipid modified oligo by flow cytometry.

**Supp figure 4**. Related to Figure 2. Additional quality control plots comparing sample multiplexing tag and SNP assignments.

**Supp figure 5**. Related to Figure 3. Additional metrics on the performance of multiplexing tag oligo reagents in mouse embryo E18.5 brain experiment.

**Supp figure 6**. Related to Figure 3. Effect of prolonged incubation on ice to oligo tag specificity.

**Supp figure 7**. Related to Figure 4. Ovarian carcinosarcoma xenograft sample multiplexing experiment.

**Supp figure 8**. Related to Figure 5. Fixed ovarian carcinosarcoma xenograft experiment.

**Supp figure 9**. Related to Figure 6. Additional metrics on the Jumpcode CRISPRclean human gRNAs on PBMCs.

**Supp figure 10**. Related to Figure 6. Performance of Jumpcode CRISPRclean human gRNAs on the Ovarian patient derived xenograft experiment.

**Supplementary Figure 1.**
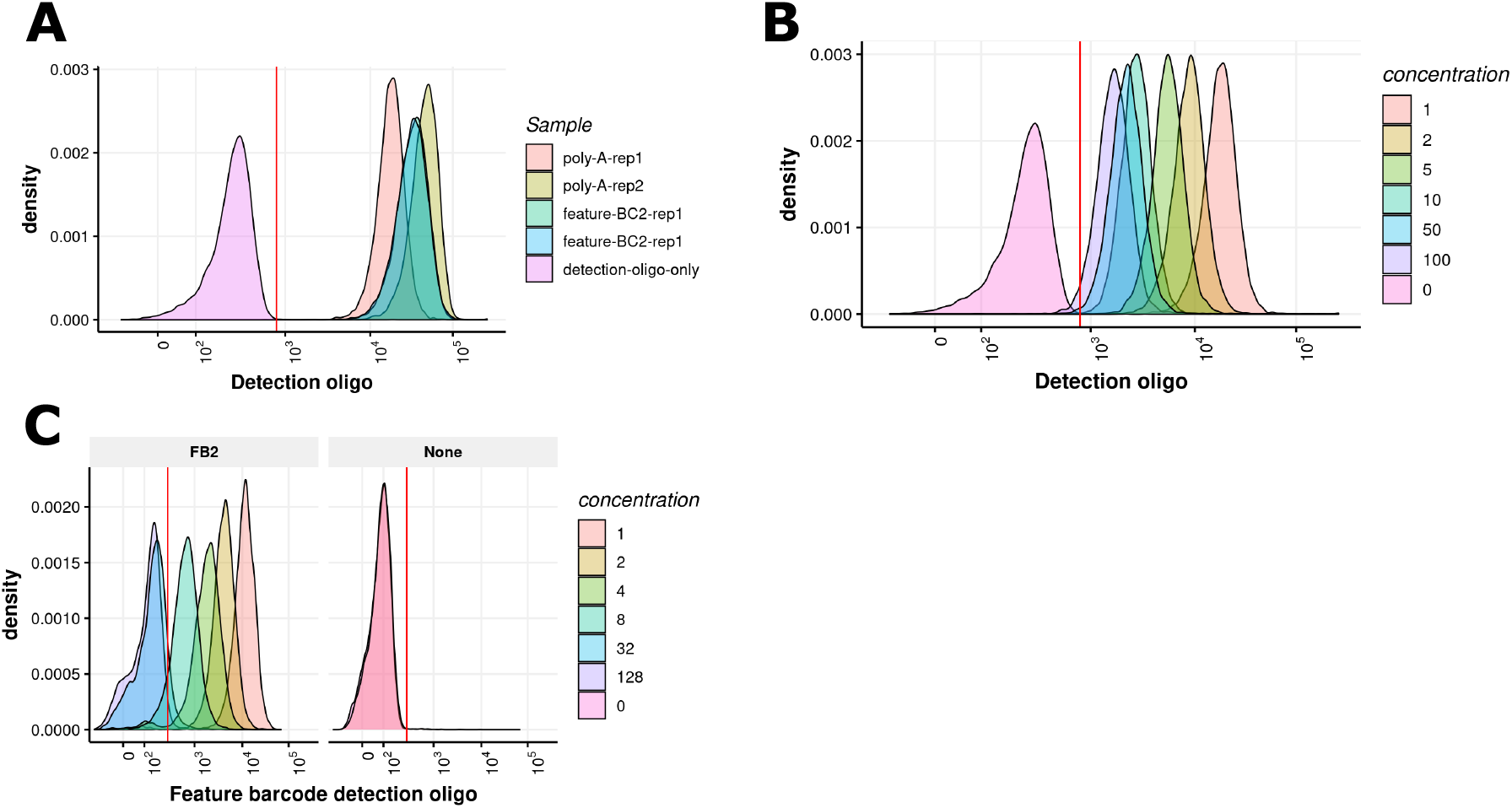
Optimisation of sample multiplexing tag oligo labelling by flow cytometry. (A) FACS comparison of original MULTI-Seq oligo poly-T capture oligo sequence with feature barcode 2 capture sequence. (B) Titration of hashtag antibody oligos. (C) Titration of MULTI-Seq LMO oligos. The detection oligo only control is shown separately for clarity.

**Supplementary Figure 2.**
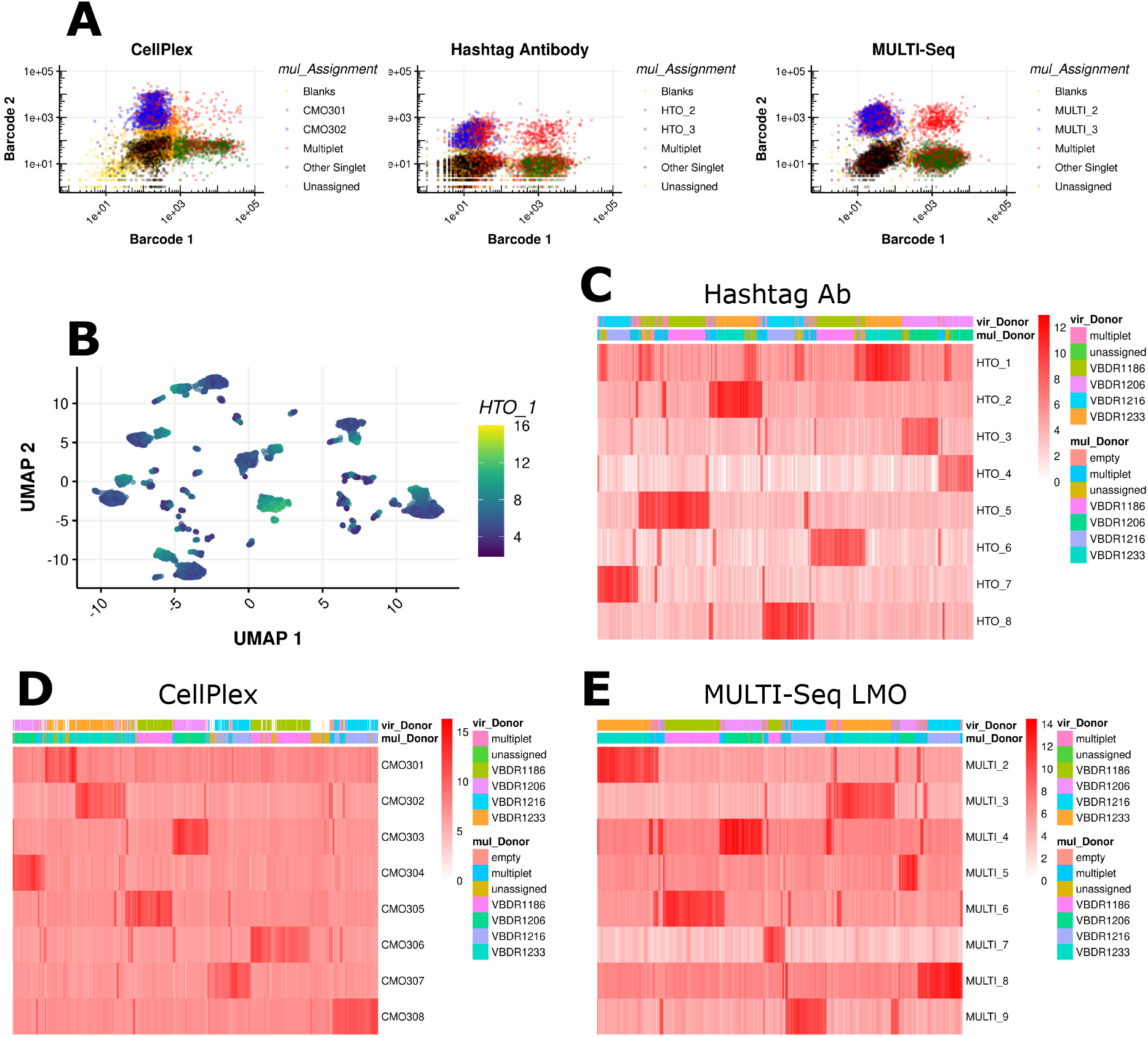
Additional metrics on the performance of sample multiplexing reagents in the PBMC experiment. (A) Parwise scatter plots of two oligo tags. Note all doublets for all tags are shown in red. (B) UMAP of hashtag antibody tag counts coloured by detection of HTO-1. (C) Heatmap of hashtag antibody tag counts. (D) Heatmap of CellPlex tag counts. (E) Heatmap of MULTI-Seq LMO tag counts.

**Supplementary Figure 3.**
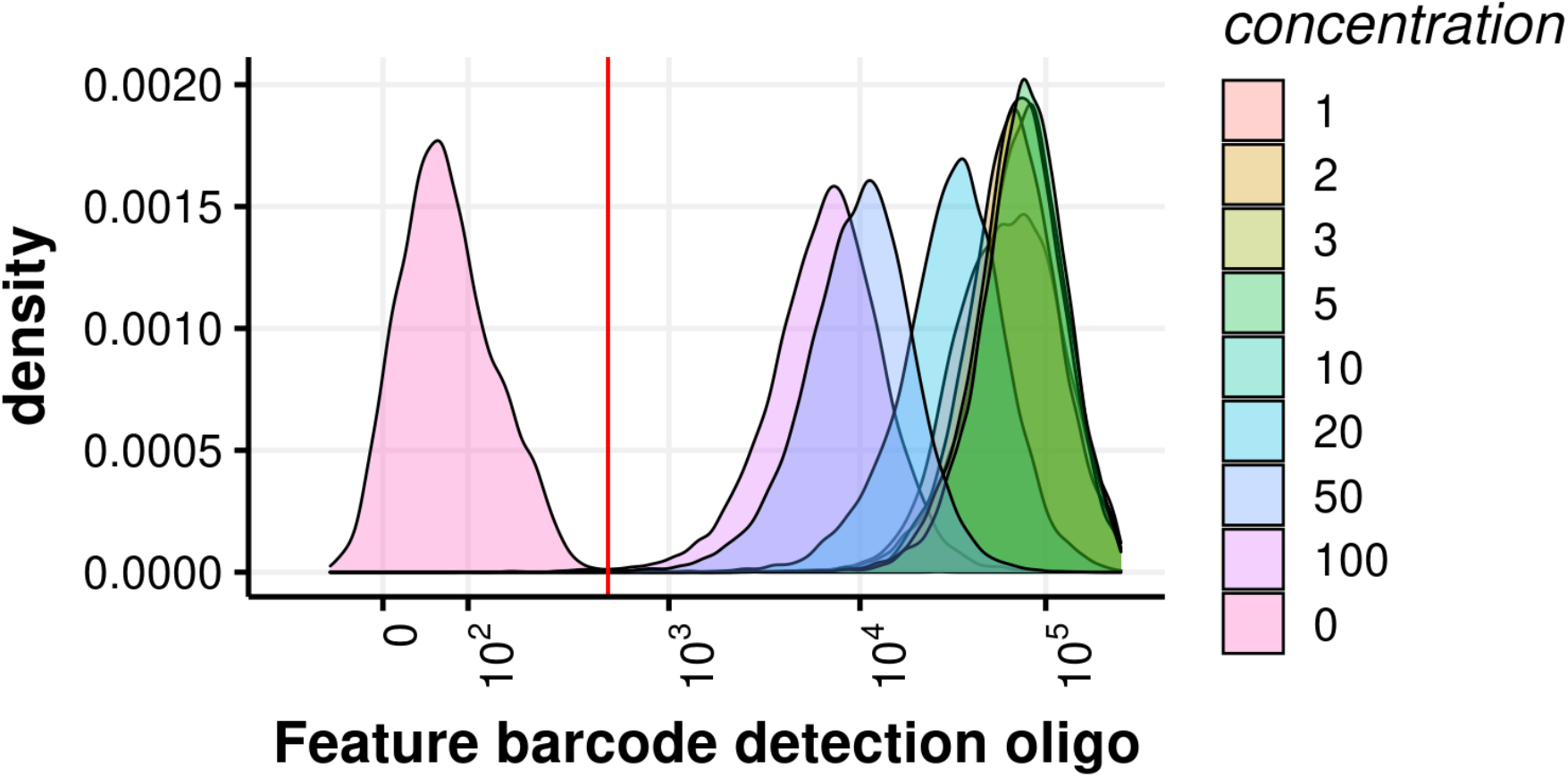
Titration of CellPlex lipid modified oligo by flow cytometry. K562 cells were labelled with a serial dilution of CellPlex. Cells were analysed by FACS after adding a feature barcode 2 fluorescent detection oligo.

**Supplementary Figure 4.**
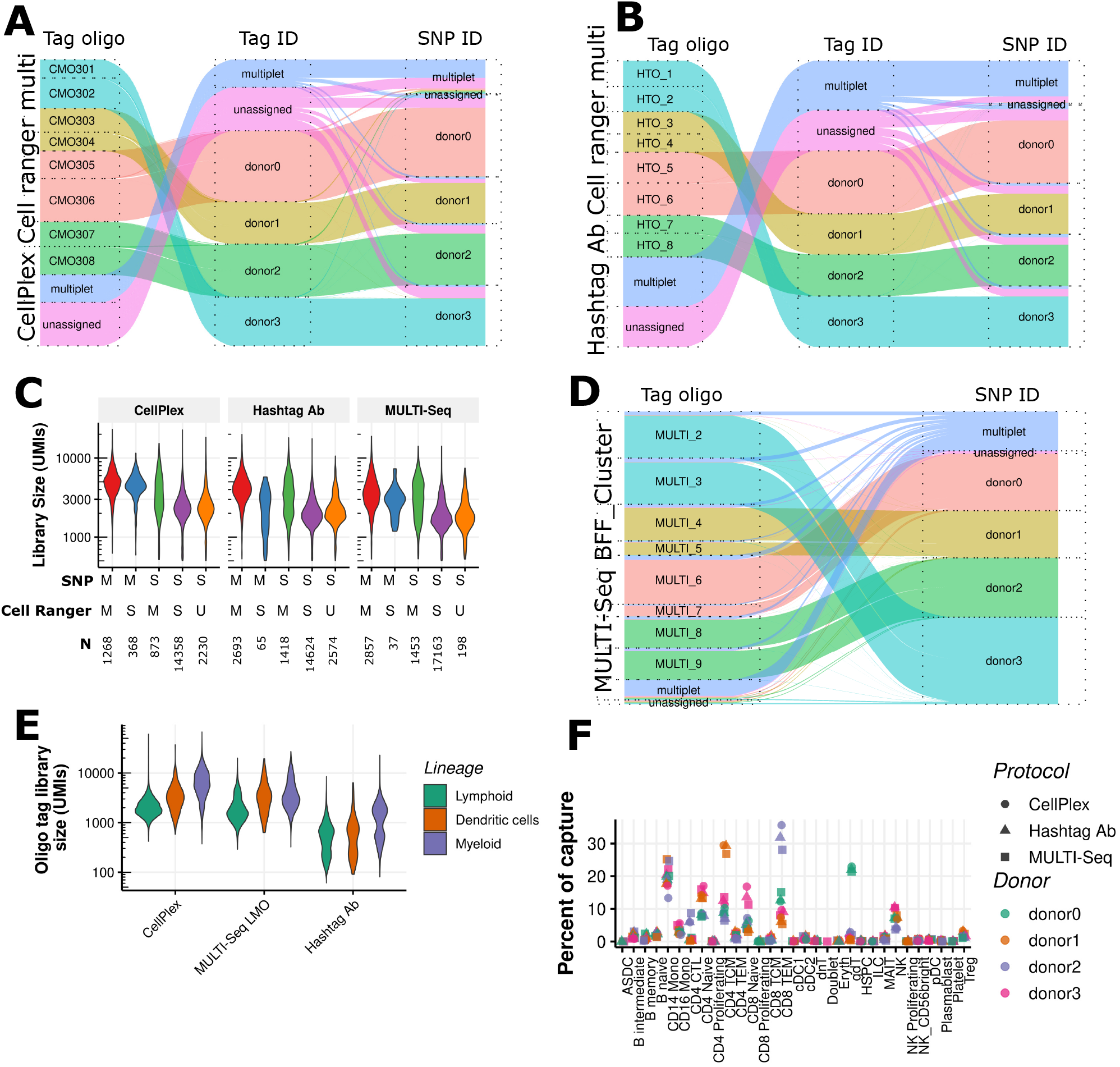
Multiplexing tag oligo and SNP assignments for PBMCs. (A) Alluvial plots for CellPlex demultiplexed with Cell Ranger multi. Left, tag oligo identity, middle donor identity based on oligo tags, right donor identity based on SNPs. (B) Alluvial plots for Total-Seq A hashtag antibody demultiplexed with Cell Ranger multi. (C) Gene expression library size comparison between oligo tags and SNP call. M = Multiplet, S = Singlet, N = number of cells in category. (D) Alluvial plots for MULTI-Seq demultiplexed with BFF Cluster. Left, tag oligo identity called by BFF Cluster, middle donor identity based on oligo tags, right donor identity based on SNPs. (E) Oligo tag library size by immune lineage. (F) Immune subset composition by donor and protocol. Automated annotation is shown with Level 2 granularity.

**Supplementary Figure 5.**
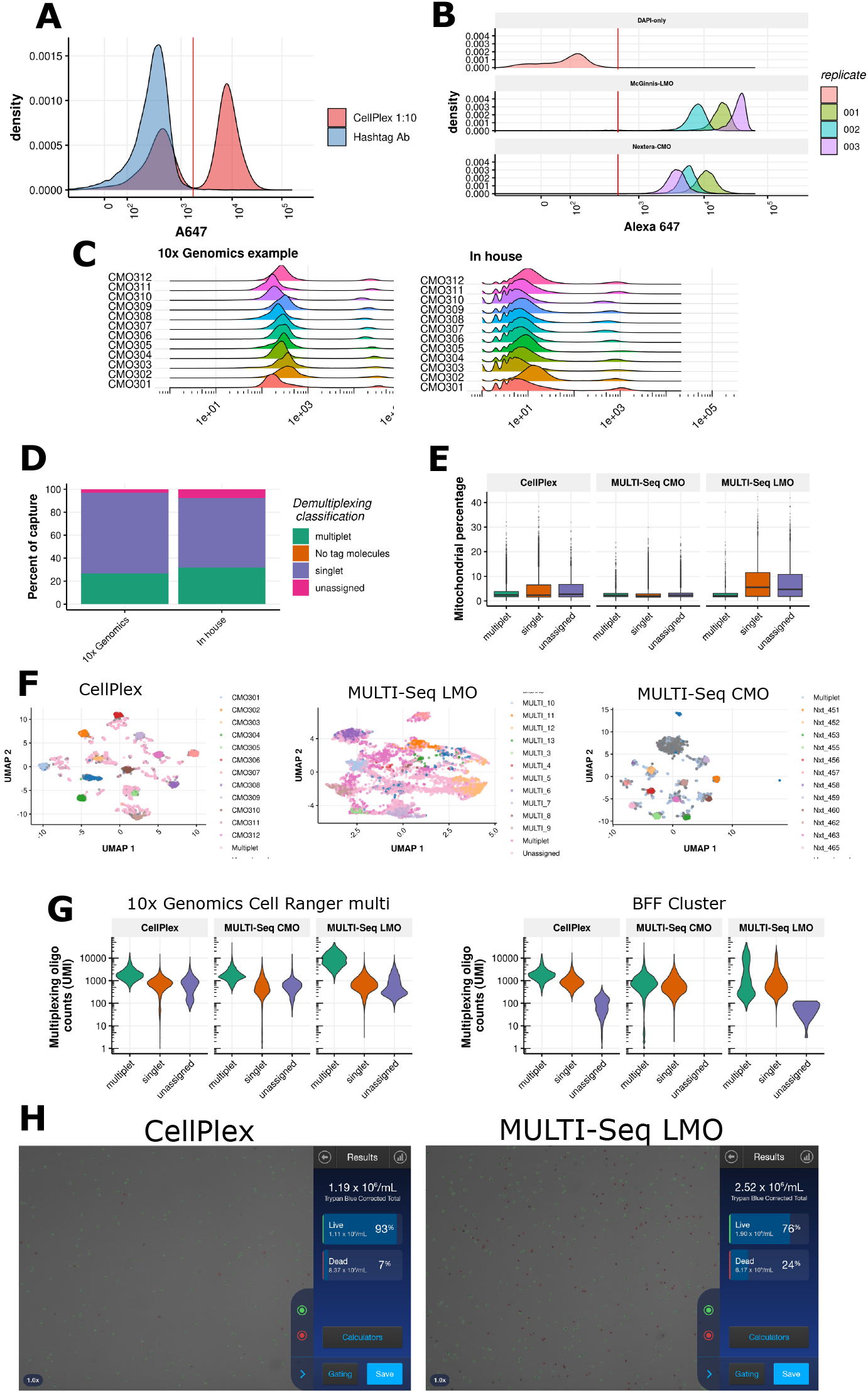
Performance of multiplexing oligo tag reagents in mouse embryo E18.5 brain experiment. (A) Comparison of mouse Total-Seq A hashtag antibody with CellPlex 1:10 dilution on mouse E18.5 brain. Signal was read out by FACS. (B) Comparison of MULTI-Seq LMO with custom MULTI-Seq cholesterol modified oligo (CMO). The Illumina small RNA read 2 sequencing handle was substituted with Nextera read 2 handle. (C) Comparison of undiluted CellPlex oligo from 10x Genomics demonstration dataset with the ten-fold dilution used in the mouse E18.5 brain experiment. (D) Droplet identity generated with Cell Ranger multi of undiluted CellPlex oligo from 10x Genomics demonstration dataset compared to the ten-fold dilution used in the mouse E18.5 brain experiment. (E) Mitochondrial count percentage for each protocol. (F) UMAP dimension reduction visualisation of multiplexing oligo tag counts for each multiplexing protocol. (G) Multiplexing oligo tag library size for each droplet assignment category and protocol. (H) Images from countess III of leftover CellPlex and MULTI-Seq LMO labelled samples taken during 10x Genomics capture.

**Supplementary Figure 6.**
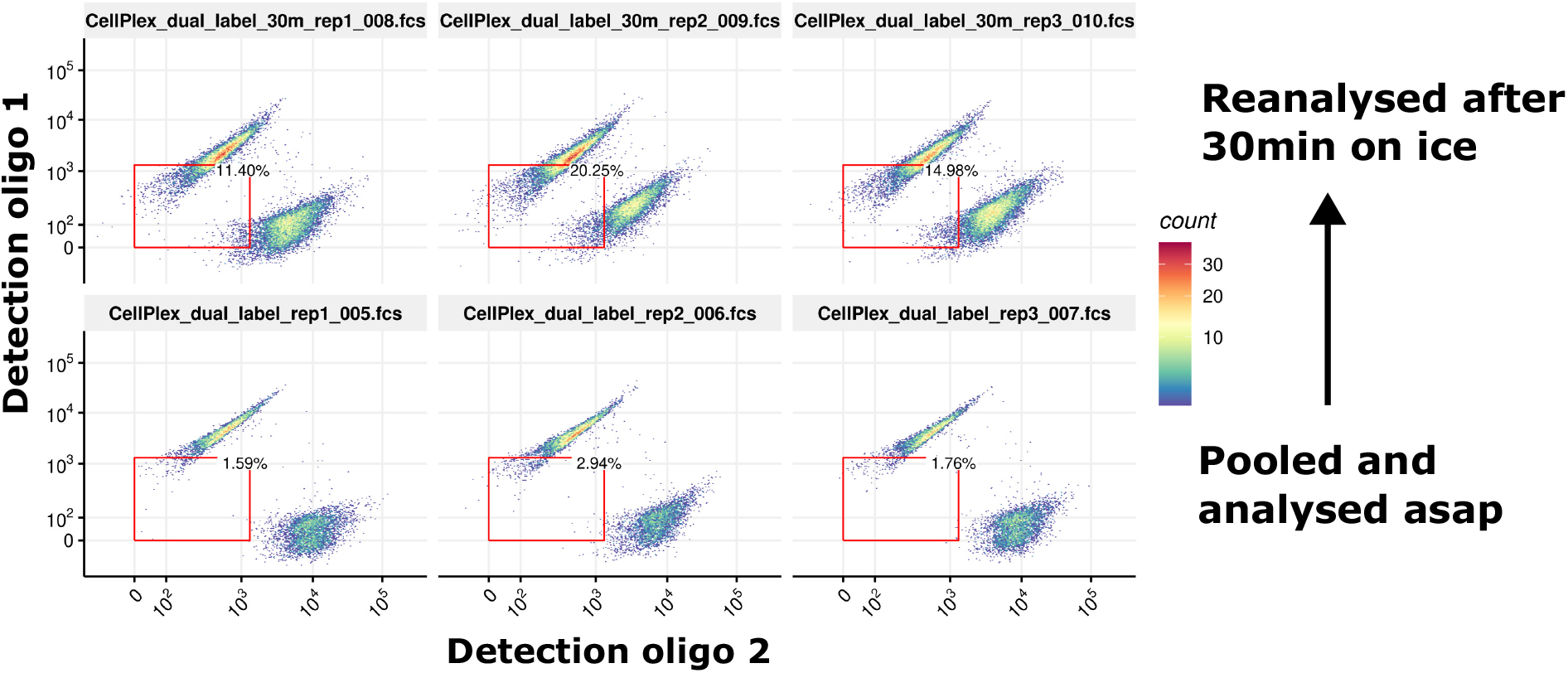
Comparison of CellPlex signal with prolonged incubation after pooling. Two distinct samples were labelled individually with CellPlex and fluorescent detection reagents and pooled immediately prior to FACS analysis (bottom) or after 30 minutes on ice (top). Experiment was performed in triplicate.

**Supplementary Figure 7.**
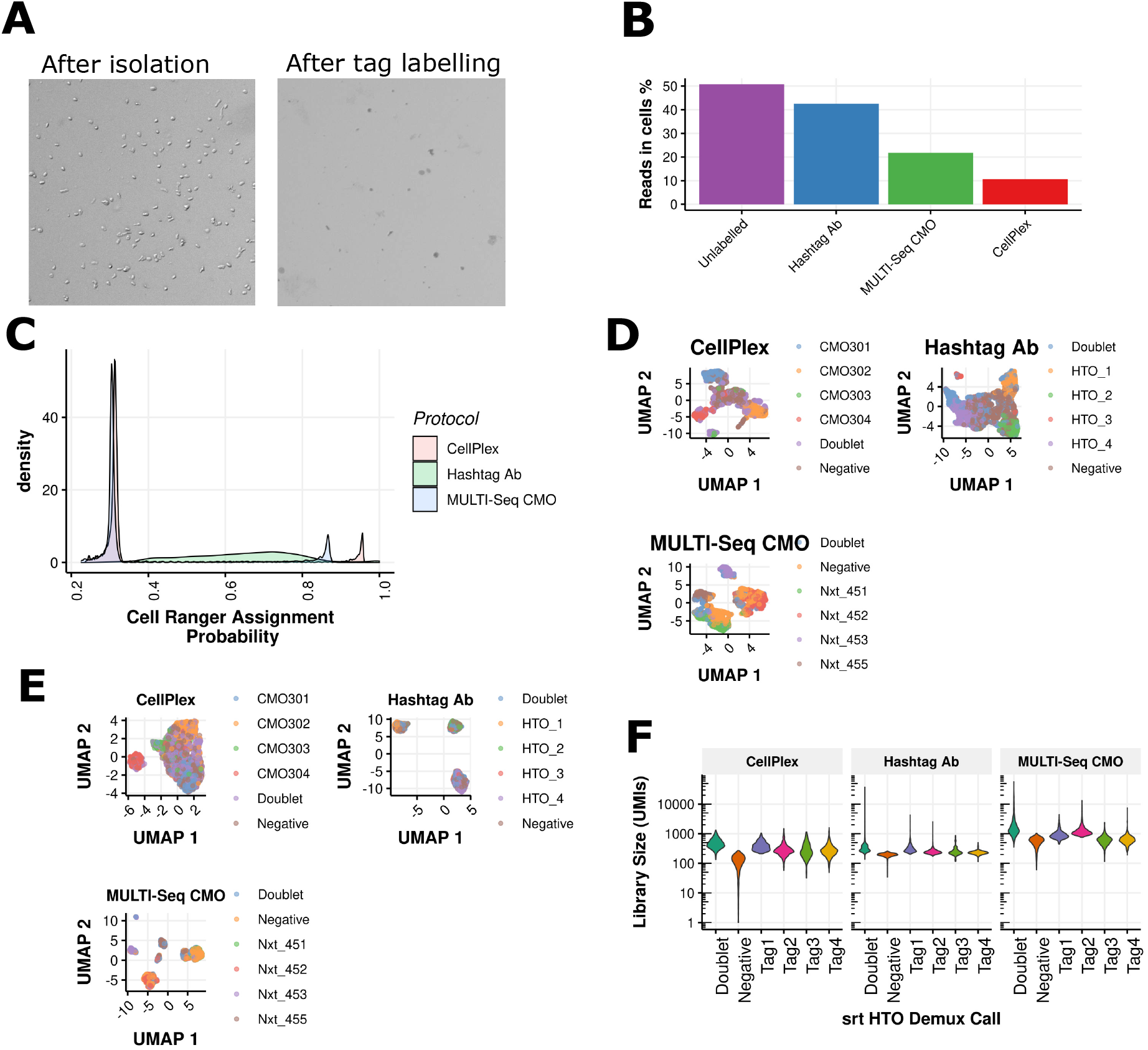
Performance of multiplexing tag oligo reagents in Ovarian carcinosarcoma xenograft experiment. (A) Countess images taken immediately after nuclei extraction with 10x Genomics kit and after multiplex oligo labelling and pooling.2x zoomed image from Countess for a total 4.5x magnification (B) Reads in cells estimate for each sample multiplexing protocol. (C) 10x Genomics Cell Ranger multi assignment scores for the three oligo tag protocols. The majority of nuclei failed the 90% threshold for assignment to a tag. (D) UMAP dimension reduction visualisation of multiplexing tag counts for each protocol. Nuclei are coloured by BFF_cluster call. (E) UMAP dimension reduction visualisation of gene expression data for each protocol. Nuclei are coloured by BFF_cluster call. (F) Oligo tag library size for distinct categorises called by Seurat HTODemux.

**Supplementary Figure 8.**
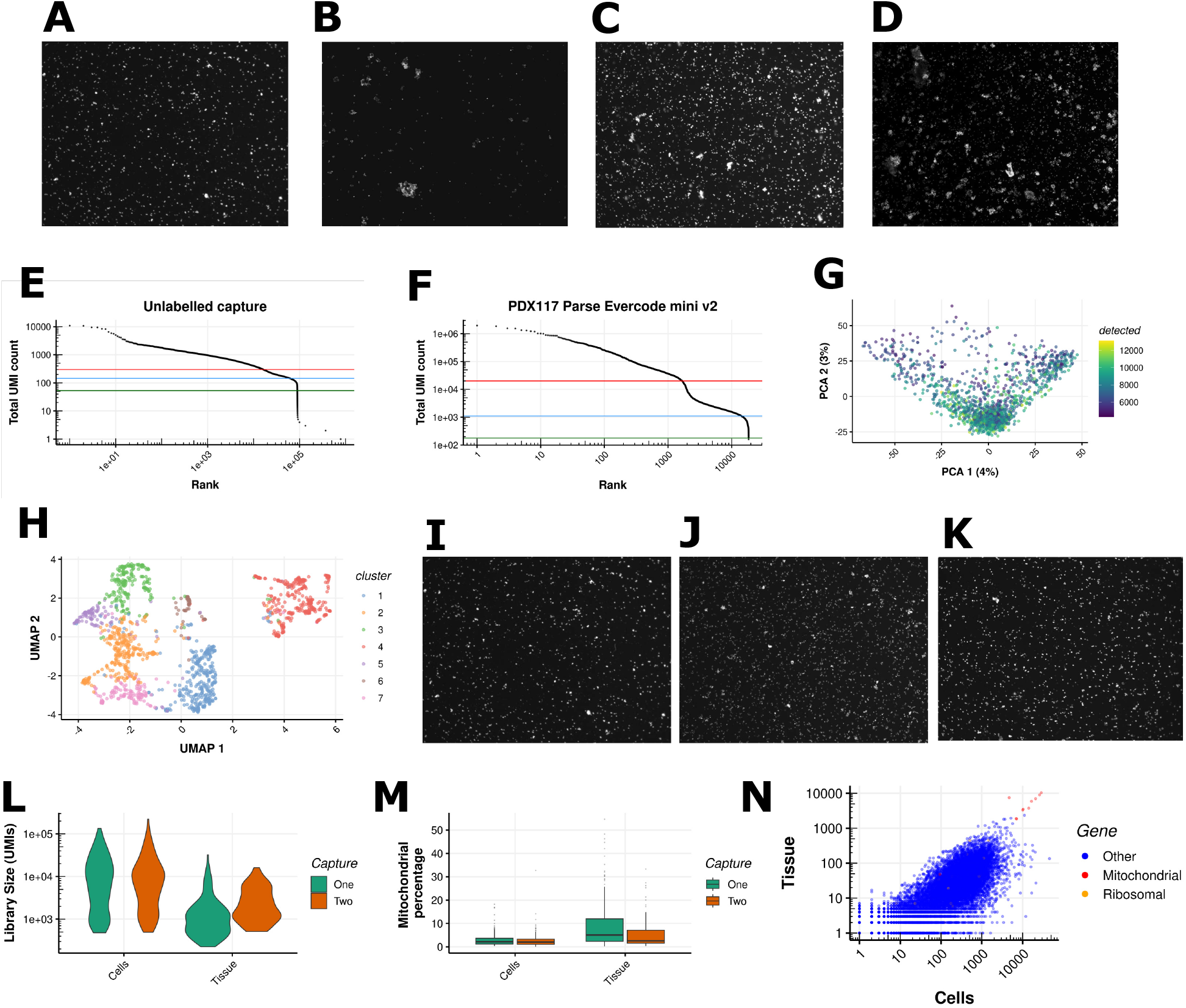
Fixed RNA-Seq ovarian carcinosarcoma xenograft experiment. (A) 10x Genomics Flex nuclei post probe hybridisation and before sample pooling. 2x zoomed image from Countess for a total 4.5x magnification. Nuclei are visualised with propidium iodide and images collected in the RFP channel. (B) Pooled 10x Genomics Flex nuclei prior to microfluidics capture. (C) Retained Parse Biosciences v2 sample prior to reverse transcription. (D) Omitted Parse Biosciences v2 sample prior to reverse transcription. (E) PCA of Parse Bioscience at full sequencing amount, coloured by number of genes detected. (F) UMAP of Parse Bioscience at full sequencing amount, coloured by cluster number. (G) 10x Genomics Flex fixed cells, post probe hybridisation and before sample pooling. (H) 10x Genomics Flex fixed tissue dissociated to cells, post probe hybridisation and before sample pooling. (I) Pooled 10x Genomics Flex experiment on cells prior to microfluidics capture.2x zoomed image from Countess for a total 4.5x magnification. Cells are visualised with propidium iodide and images collected in the RFP channel. (J) Comparison of mitochondrial gene percentage from 10x Genomics Flex experiment in cells and tissues. (K) Comparison of library size from 10x Genomics Flex experiment in cells and tissues. (L) Comparison of number of genes detected from 10x Genomics Flex experiment in cells and tissues. (M) Gene expression comparison of fixed cells versus fixed tissue for 10x Genomics Flex experiment. Each dot is the sum of counts across all single cells for a gene.

**Supplementary Figure 9.**
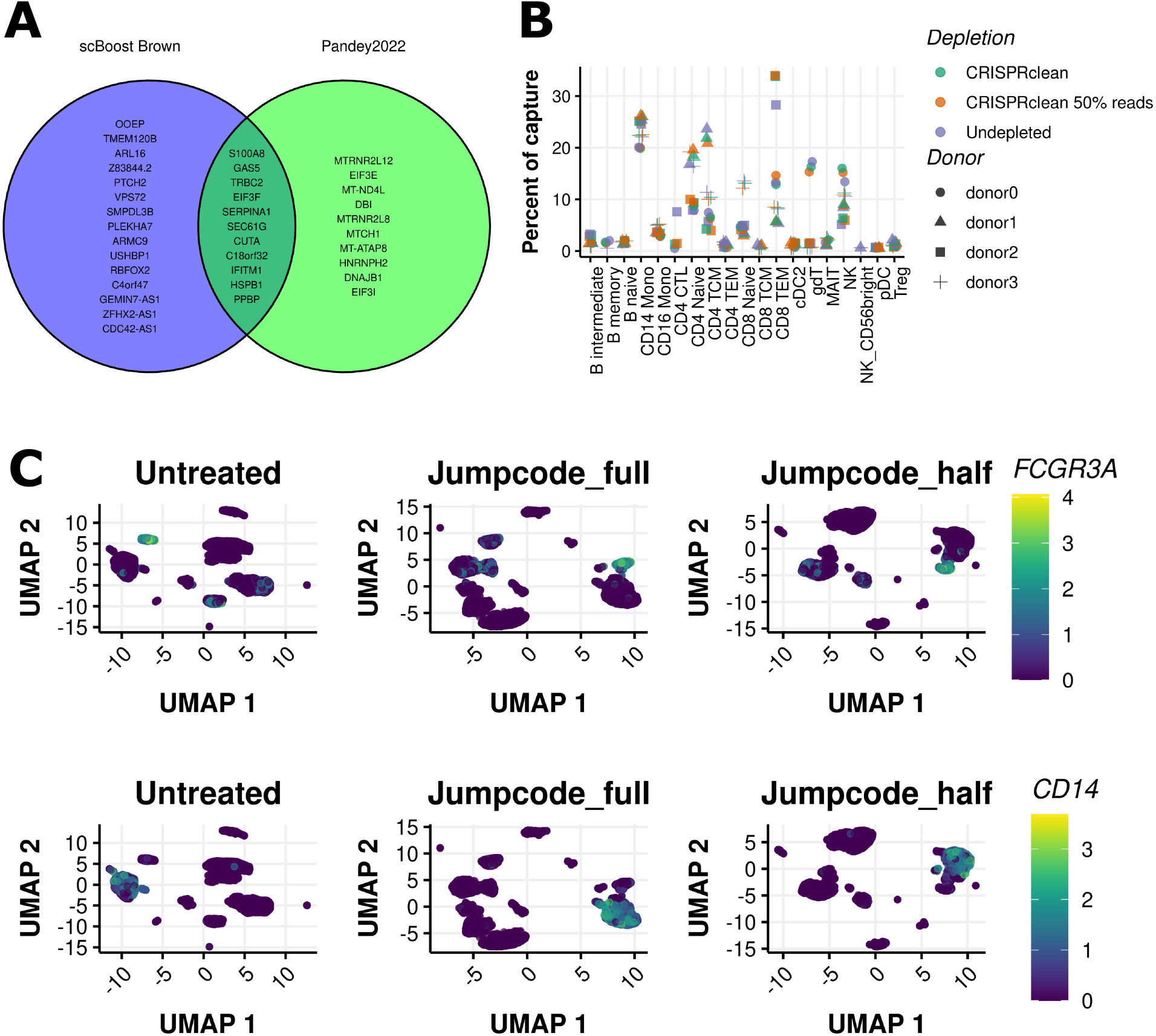
Additional metrics on the Jumpcode CRISPRclean human gRNAs on PBMCs. (A) Summary of immune subsets recovered from untreated, CRISPRclean treated, and downsampled CRISPRclean treated gene expression data. (B) Venn diagram listing off target genes differentially decreased in PBMC experiment with the list of genes from Pandey et al., 2022. (C) Expression of *FCGR3A*, (CD16, non-classical monocytes) and *CD14* (classical monocytes) in untreated, CRISPRclean treated and 50% downsampled CRISPRclean treated libraries.

**Supplementary Figure 10.**
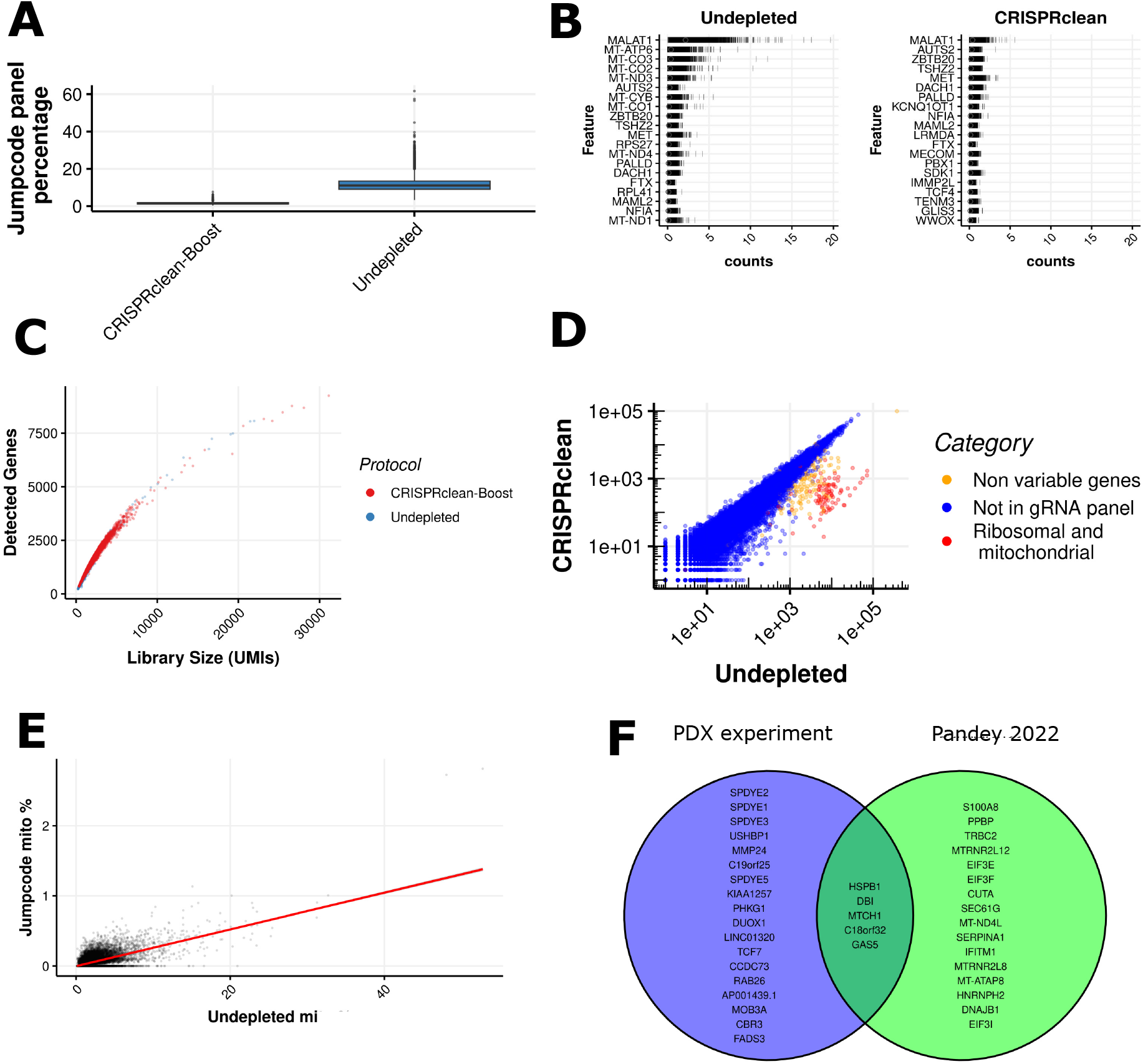
Performance of Jumpcode CRISPRclean human gRNAs on the Ovarian patient derived xenograft experiment. (A)Relationship between library size and number of detected genes per cell. (B) Top 20 highly expressed genes in CRISPRclean treated and untreated libraries. Genes prefixed as “RP” or “MT-” represent ribosomal or mitochondrial genes, respectively. (C) Relationship between library size and number of detected genes per cell. (D) Gene expression comparison, each dot represents the sum of counts across all single cells for a gene. (E) Correlation of mitochondrial gene percentages. The trend line is a linear fit. (F) Venn diagram listing off target genes differentially decreased in Ovarian carcinosarcoma PDX experiment.

